# Tissue-specific restriction of TE-derived regulatory elements safeguards cell-type identity

**DOI:** 10.1101/2025.05.13.653700

**Authors:** Danica Milovanović, Julien Duc, Wayo Matsushima, Romain Hamelin, Evarist Planet, Sandra Offner, Olga Rosspopoff, Didier Trono

## Abstract

Transposable elements (TEs) have extensively reshaped the *cis*-regulatory landscape of mammalian genomes, yet the mechanisms that govern their context-specific activity remain incompletely understood. KRAB zinc finger proteins (KZFPs), a large family of transcription factors specialized in TE recognition, are known repressors of TE-derived regulatory activity through TRIM28-mediated H3K9me3 deposition. Here, we expand this paradigm by uncovering noncanonical relationships between TEs and KZFPs. By generating a comprehensive epigenomic map of KZFP-bound TEs, we find that the regulatory activity of ancient mammalian L2/MIR elements is broadly delineated by KZFP binding patterns, despite low H3K9me3 enrichment. We further dissect this relationship by investigating the function of ZNF436, a non-canonical KZFP, highly expressed during *in vivo* human fetal heart development. Using loss-of-function approaches, we show that ZNF436 preserves cardiomyocyte function by promoting cardiac gene expression while restricting the activation of alternative lineage programs. Mechanistically, ZNF436 recruits specialized SWI/SNF chromatin remodeling complexes to limit the accessibility of L2/MIR-derived enhancers, many of which are active in non-cardiac tissues. These findings reveal a noncanonical, TRIM28-independent role for KZFPs in shaping cell-type-specific regulatory landscapes and emphasize the importance of repressing alternative regulatory programs alongside activating lineage-specific ones to safeguard cell identity.

## INTRODUCTION

The specification of cell fate and identity is governed by a complex interplay of epigenetic mechanisms and chromatin regulation, ensuring the timely activation of tissue-specific genes and the repression, or poising, of alternative developmental programs (Barrero, et al. 2010). These gene expression programs are orchestrated by intricate regulatory networks that emerge from dynamic interactions between transcription factors (TFs), their associated *cis*-regulatory elements (CREs), and the broader chromatin landscape in which they operate (Carroll, 2008; Fillot and Mazza, 2025). Importantly, both CREs and TF binding sites have undergone extensive evolutionary diversification across vertebrate lineages, with fewer than 5% of human CREs conserved beyond mammals (Osmanski et al., 2023; Andrews et al., 2023). This rapid turnover raises the question of whether, and how, such dynamic regulatory landscapes shape chromatin states and gene expression in a cell-type-specific manner.

To address this question, we focus on the interplay between transposable elements (TEs) and Krüppel-associated box (KRAB) zinc finger proteins (KZFPs), two rapidly evolving genomic components that contribute to mammalian species-specific regulatory layers. TEs make up over half of the human genome (Lander, et al., 2001; Andrews, et al., 2023) and were historically considered parasitic due to their ability to mobilize within genomes and induce ectopic recombination events (Ohno, 1970; Orgel and Crick, 1980). Yet, fewer than 0.05% of TEs remain mobile, with the vast majority rendered static and degenerated by accumulated mutations (Mills, et al., 2007; Matsushima, et al., 2024). These TE-derived sequences have been extensively repurposed by the host in a process known as TE co-option or exaptation. As envisioned by Barbara McClintock’s concept of “controlling elements” (McClintock, 1950), co-opted TEs act as enhancers, insulators, TF binding sites, and sources of novel promoters and exons (Chuong et al., 2017). Critically, because TE subfamilies differ markedly between species, their contributions to gene regulation are often species- or lineage-specific, positioning TEs as major drivers of regulatory innovation. Recent estimates suggest that ∼25% of human candidate CREs derive from TEs (Du, et al., 2024), and that 20% of all human TF binding sites reflect primate-specific innovations linked to TE insertions (Andrews, et al., 2023). The widespread genomic distribution of TEs and their inherent sequence diversity make them ideal platforms for coordinated, context-dependent gene regulation. Indeed, the regulatory activity of TE-derived sequences is highly dynamic, varying across developmental stages and cell types, including genome activation, imprinting, gastrulation, tissue specification, and brain development (Bourque, et al. 2018; Pontis, et al. 2019; Playfoot, et al. 2021; Gassler, et al. 2022; Iouranova, et al. 2022). The pervasive co-option of TEs into gene regulatory networks creates a biological trade-off between limiting their potentially deleterious effects and exploiting their regulatory potential. This balance is largely mediated by KZFPs, the largest TF family in mammals (Imbeault, et al., 2017). In the human genome, the KZFP gene family comprises nearly 400 members, encoding proteins that exhibit exceptional specificity toward TE-derived sequences (de Tribolet-Hardy, et al., 2023). As a result, the repertoire of KZFPs is also highly species-specific, with individual KZFPs often regulating one or few related TE subfamilies (Imbeault, et al., 2017; Helleboid, et al., 2019; de Tribolet-Hardy, et al., 2023). KZFPs typically contain an N-terminal KRAB domain, which interacts with TRIM28/KAP1 to induce H3K9me3-mediated repression, and a C-terminal C2H2 zinc finger array that confers sequence-specific DNA binding (Rosspopoff and Trono, 2023). These properties make KZFPs well-suited to commission context-dependent roles for TE-derived elements, as illustrated by their H3K9me3-mediated attenuation of TE enhancer activity (Pontis, et al., 2019; Iouranova, et al. 2022; De Franco, et al., 2023). However, around 10% of human KZFPs harbor variant KRAB domains that lack TRIM28 interaction, some of which contain accessory domains that enable functional diversification (Rosspopoff, et al., 2025; Matsushima, et al., 2025), also correlated by the very distinct protein interactomes of these non-canonical KZFPs (Helleboid, et al., 2019). Yet, how KZFPs function outside of H3K9me3 deposition and whether they guide cell-type-specific regulation of TE-derived CREs remains poorly understood.

In this study, we explored non-canonical and context-dependent roles of KZFPs in regulating TE-derived elements. We identified ZNF436, a KZFP endowed with a variant KRAB domain, as a regulator of L2/MIR-derived CREs. Using an *in vitro* model of cardiomyocyte differentiation, we show that ZNF436 plays a dual role by maintaining cardiac gene expression and limiting the ectopic activation of genes associated with alternative lineages. Mechanistically, ZNF436 functions independently of the canonical TRIM28 repression axis and instead selectively engages with a specialized SWI/SNF remodeling complex to restrict the activation of L2/MIR-derived CREs, typically active in non-cardiac tissues. This reveals a novel mechanism by which KZFPs achieve enhancer selectivity and transcriptional precision in a specific lineage, extending the classical TE/KZFP paradigm into tissue-specific enhancer modulation during development.

## RESULTS

### KZFP binding delineates TEs endowed with CRE activity

To refine our understanding of TE regulatory behavior, we systematically integrated KZFP binding and associated TEs with epigenetic regulatory features using publicly available datasets (Fig. 1A) (Schmitges, et al. 2016; Imbeault, et al. 2017; The ENCODE Project Consortium, et al. 2020; de Tribolet-Hardy, et al. 2023). First, we stratified TE subfamilies according to their statistically significant association with individual KZFPs. We then quantified the enrichment for six histone marks of KZFP-stratified subfamilies across up to 82 tissues (Fig. 1A, see methods). This approach described each TE subfamily with a vectorial epigenomic signature, defined by values summarizing different combinations of histone mark enrichment across tissues together with the KZFP binding. To explore the relationships among TE subfamilies based on these signatures, we visualized them using UMAP dimensionality reduction (McInnes, et al. 2018). Strikingly, despite this method being agnostic to the shared evolutionary histories of TEs, the spatial organization in UMAP space mirrored known TE class and family relationships (Fig. 1B, and fig. S1A). This pattern aligns with the target sequence specificity of KZFPs, which often bind one or several evolutionary related TE subfamilies (de Tribolet-Hardy, et al., 2023). For instance, the relatively young LINE-1 subfamilies (∼20 million years old on average), which are known targets of multiple KZFPs (Forey, et al., 2025), formed a distinct group (L1PA1-L1PA16, black arrow, Fig. 1B). Interestingly, we also observed that older TEs from distinct families, such as the LINE/L2s and SINE/MIRs (collectively referred to as L2/MIR), appeared as nearest neighbors in the UMAP space (Fig. 1, B and C). Despite differences in sequence structure, this proximity suggested functional convergence, possibly driven by shared sequence fragments. Supporting this hypothesis, the non-autonomous MIR elements are known to have hijacked L2-derived sequences for their propagation approximately 160 million years ago (Fig. 1C) (Lander, et al. 2001; Kramerov and Vassetzky, 2011). Supporting the specificity of this observation, other nonautonomous elements such as SINE/Alu were located far from their LINE1 partners (Fig. 1B). This was consistent with their regulation by distinct sets of KZFPs (Deininger, 2011; de Tribolet-Hardy, et al. 2023) and strongly suggested that the noted proximity of L2 and MIR on the UMAP space was not serendipitous.

**Fig. 1.**
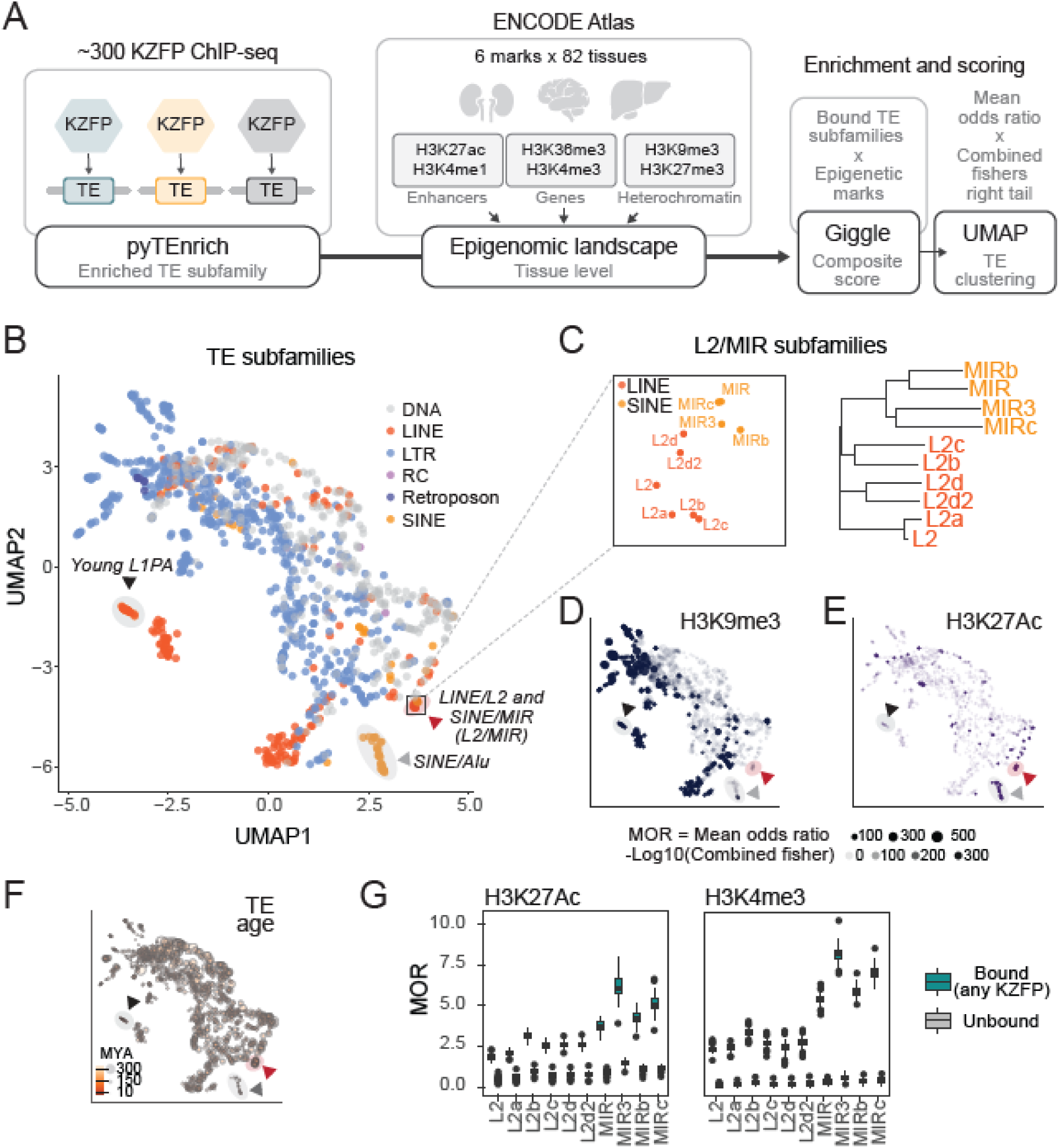
KZFP binding stratifies TE subfamilies enriched in specific chromatin states. (**A**) Schematic overview of the data analysis pipeline used to compute GIGGLE scores for histone mark enrichment across tissues for KZFP-stratified TE subfamilies. (**B**) UMAP representation of GIGGLE scores for each TE subfamily, colored by TE class. Black, grey, and red arrows indicate well-defined groups of LINE1/L1PA, SINE/Alu, and L2/MIR subfamilies, respectively. RC, Rolling Circle TE. (**C**) Phylogenetic tree of L2/MIR subfamilies based on the multiple sequence alignment of Dfam consensus sequences. (**D** and **E**) UMAP projection of the mean odds ratio (MOR) enrichment score for H3K9me3 (**D**) and H3K27ac (E) histone marks for each TE subfamily. (**F**) UMAP projection of TE subfamily ages, based on manual curation from Dfam database. (**G**) MOR for H3K27ac and H3K4me3 histone marks for L2/MIR subfamilies across tissues, grouped by KZFP-bound and unbound integrants. All KZFP-bound integrants were significantly enriched compared to unbound ones (p-value < 0.001). P-values were computed using a paired t-test.

Building on these observations, we next examined how specific histone marks associate with KZFP-stratified TE subfamilies across tissues. As expected for young TE subfamilies subjected to KZFP-mediated repression, such as L1PAs, strong association with H3K9me3 was observed across tissues (Fig. 1D). In contrast, older L2/MIR elements displayed a more active chromatin signature, with marked enrichments for H3K27ac and H3K4me3 (Fig. 1E, and fig. S1B), consistent with their known roles as enhancers and promoters (Jjingo, et al. 2014; Cao, et al. 2019; Roller, et al. 2021). Thus, the distinction between repressive and active chromatin states correlated broadly with the evolutionary age of TE subfamilies (Fig. 1F). Although this age-related pattern was previously reported (de Tribolet-Hardy, et al. 2023; Hyacinthe and Bourque, 2024), such trend persisted in our analysis of KZFP-stratified TE subfamilies, including older subfamilies that were not broadly associated to a repressed chromatin state. This supports a model in which KZFPs can not only repress TEs through heterochromatin deposition, but also associate with broadly active, co-opted CREs. To further test this model, we compared the average histone mark signals across tissues for L2/MIR integrants bound by at least one KZFP versus unbound ones. While KZFP-bound L2/MIRs displayed higher enrichment for all histone marks, this tendency was more pronounced for active chromatin states typically associated with gene proximal and distal regulatory regions across tissues (e.g. H3K4me3 and H3K27Ac, Fig. 1G and fig. S1, C and D). These results establish that the systematic integration of KZFP binding profiles allows to delineate TE integrants with regulatory potential within subfamilies. Finally, the separation of L2/MIRs from other TE subfamilies with similar chromatin signatures in the UMAP space (Fig. 1E) suggested that KZFP binding might contribute to their observed spatial segregation, pointing to a regulatory role distinct from classical repression.

### A strong L2/MIR-binder, ZNF436 evolves under purifying selection

To investigate which KZFPs delineate the CRE activity of L2/MIRs, we focused on KZFPs that were statistically enriched over one or several L2/MIR subfamilies. We identified 30 KZFPs, five of which recognize L2/MIRs as their primary TE targets (ZNF436, ZNF3, ZKSCAN8, ZKSCAN1, and ZNF662, Fig. 2A). Except for the ∼100 myo ZNF662, the remaining four KZFPs emerged concomitantly with transposition-competent L2/MIRs over 160 million years ago. Strikingly, all four KZFPs carry a so-called variant KRAB (vKRAB) domain lacking H3K9me3-mediated repressor activity, a feature shared by only ∼10% of all KZFPs (Tycko, et al. 2020; Rosspopoff and Trono, 2023). Therefore, these KZFPs stood out as promising candidates regulating L2/MIR-derived CRE activity, since it is characterized by an active chromatin state and a relatively low H3K9me3 enrichment (Fig. 1G, and fig. S1C). Amongst them, ZNF436 further stood out by targeting 9 out of 10 subfamilies of L2/MIRs, with the highest fold change enrichment values for L2b and L2c (FCE of 14.6 and 10.6, respectively, Fig. 2A, and fig. S2A).

**Fig. 2.**
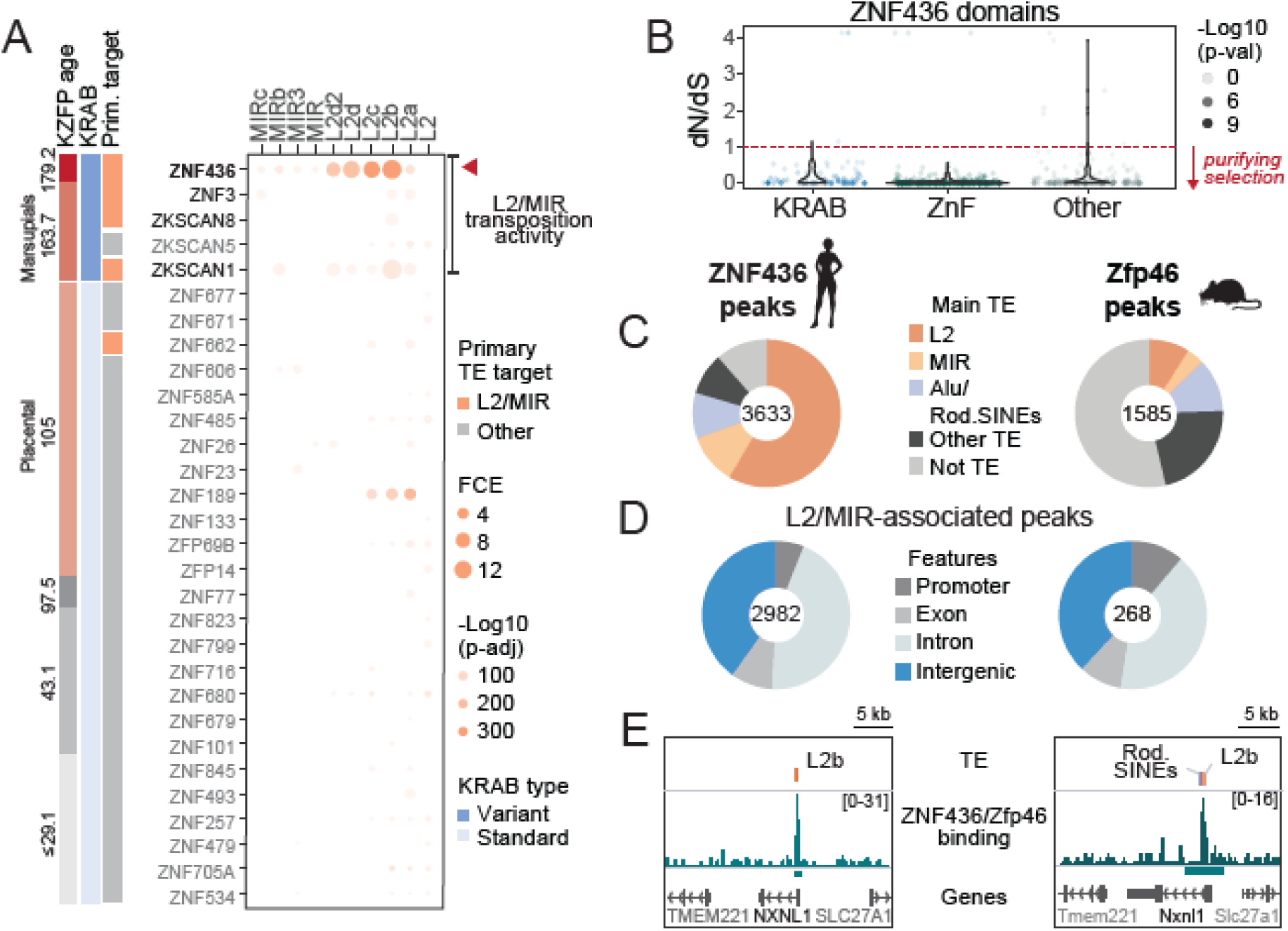
ZNF436 is a conserved KZFP with a strong binding affinity for L2/MIR integrants. (**A**) Enrichment of KZFP binding over L2/MIR subfamilies. The primary TE target indicates whether any of the L2/MIR subfamilies is the top-enriched TE subfamily for a given KZFP. Estimates of KZFP age and presence of standard/variant KRAB domain were taken from Helleboid, et al (2019). The period of L2/MIR transposition activity was obtained from Dfam. KZFPs included show significant enrichment for at least one of the L2/MIR subfamilies. Enrichment values for ZNF436 for L2b and L2c were capped at a p-adjusted of 1e-300. P-values were computed using a binomial test, adjusted for multiple testing using the Benjamini-Hochberg method. (**B**) Ratio of non-synonymous (dN) to synonymous substitutions (dS) computed for each amino acid of the ZNF436 protein sequence, grouped based on domain annotation. Values below the dotted red line (dN/dS < 1) indicate amino acids evolving under purifying selection. (**C**) Proportions of ZNF436 and Zfp46 (ZNF436 mouse ortholog) peaks associated with a specific TE group. (**D**) Proportions of ZNF436 and Zfp46 peaks overlapping L2/MIR elements, associated with the indicated genomic features. (**E**) Example genomic regions with conserved ZNF436/Zfp46 binding in the promoter regions of *NXNL1/Nxnl1* in both human and mouse genomes. Rod. SINE, rodent-specific SINE elements.

As the strongest pan-L2/MIR binder, we next addressed the functional conservation of ZNF436 by analyzing its sequence across mammals. Although some vKRAB domains may be degenerating (Emerson and Thomas, 2011), the vKRAB domain of ZNF436 is highly conserved across mammalian species (dN/dS <1, Fig. 2B, and fig. S2B), suggesting that it evolves under strong purifying selection. The same pattern was observed for its zinc finger array, with all 12 zinc fingers displaying strictly identical DNA-contacting amino acids in mammals (dN/dS = 0; Fig. 2B, and fig. S2B). Thus, it is likely that ZNF436 retained its vKRAB-mediated function and DNA binding specificity since its emergence. Supporting this, ChIP-seq data of Zfp46 (Wolf, et al. 2020), the mouse ortholog of ZNF436, showcased the enrichment for MIR-derived sequences, as well as preferential binding to L2b and L2c elements (Fig. 2C, and fig. S2C). The L2/MIR-associated binding shows a similar association with distinct genomic features in both species (Fig. 2D). Interestingly, both ZNF436 and Zfp46 showed a marked enrichment for clade-specific SINE elements (SINE/Alu in human and various rodent-specific SINEs in mouse), highlighting that a secondary wave of TE retrotransposition invaded their binding sites in respective species (Fig. 2C, and fig. S2D). For instance, the binding of ZNF436 and Zfp46 was detected at *NXNL1/Nxnl1* promoter region in both human and mouse (Fig. 2E). In humans, this region could be traced back to the insertion of an L2b element, whereas in mice, multiple rodent-specific SINEs transposed into the same locus, leaving only a small flanking L2b fragment (Fig. 2E). Promoter sequence differences in this gene have been shown to affect expression patterns in various cell types (Lambard, et al., 2010). This implies that species-specific variations are likely to occur in the regulatory output of ZNF436 and Zfp46 at specific genomic loci in their respective genome, consistent with a continuous reshaping of CRE landscapes *via* TEs insertions (Andrews, et al., 2023). Taken together, we identified a highly conserved KZFP, ZNF436, that exhibits a strong binding preference toward L2/MIR-derived sequences. Its highly conserved vKRAB domain suggests that ZNF436 regulates L2/MIR elements by mechanisms different from heterochromatin deposition.

### ZNF436 is necessary for proper cardiomyocyte contractility

To explore how ZNF436 binding could contribute to tissue-specific regulation of L2/MIRs, we first analyzed *ZNF436* expression pattern. Given that KZFP-mediated control of TE-embedded CREs plays critical roles during embryonic development (Rowe, et al. 2012; Yang, et al. 2017; Pontis, et al. 2019; De Franco, et al. 2023), we focused on gene expression datasets encompassing human *in vivo* embryonic and fetal development (Cardoso-Moreira, et al. 2019; Cao, et al. 2020). Analysis of these datasets revealed that *ZNF436* expression was highest in muscle and heart fetal tissues (Fig. 3A) (Cao, et al. 2020), and strongest in the prenatal stages, concomitant with well-known cardiac development-related genes such as *GATA6*, *TBX20*, and *TNNI1* (Fig. 3B) (Cardoso-Moreira, et al. 2019). Consistently, single-cell expression of *ZNF436* was the highest in endothelial cells and cardiomyocytes of the human fetal heart (fig. S3A) (Cao, et al. 2020). In agreement with these *in vivo* observations, ZNF436 was strongly upregulated during *in vitro* differentiation of human embryonic stem cells (hESCs) into cardiomyocytes (fig. S3B) (Zhang, et al. 2019), pointing towards a regulatory role for ZNF436 in fetal-like cardiomyocytes. Building on these observations, we assessed the transcriptomic impact of ZNF436 depletion using a CRISPRi approach in hESCs-derived ventricular-like cardiomyocytes (vCMs) (Liang, et al. 2018) at day 15 of differentiation (d15, Fig. 3C), corresponding to the peak of *ZNF436* expression (fig. S3B). ZNF436 knock-down (KD) induced widespread transcriptional changes in vCMs, with 398 upregulated and 527 downregulated genes (Fig. 3D). Notably, the expression of key vCM differentiation markers (*TNNT2, TBX5, GATA4, MEF2C*) was only modestly reduced following ZNF436 KD, and most changes did not reach statistical significance (Fig. 3D, and fig. S3C). Therefore, we concluded that the extent of transcriptomic changes observed upon ZNF436 KD does not primarily result from defects in vCM cell-fate commitment. Gene Ontology enrichment analysis on misregulated genes revealed only significantly enriched biological processes for the downregulated genes, mainly terms related to muscle development and myofibril assembly (fig. S3D). Gene set enrichment analysis (GSEA) further confirmed a strong depletion of myofibril assembly genes (Fig. 3E), which encode proteins forming functional protein interactions within contractile sarcomeres (Fig. 3F). Therefore, we asked whether the contractility of ZNF436-depleted vCMs was affected by deriving the beating frequency of d15 cardiomyocyte monolayers using particle image velocimetry (PIV) (Kumar, et al. 2019) (Fig. 3G, and fig. S3E, see methods). Strikingly, ZNF436 KD reduced by half the mean dominant beating frequency (1.77 Hz) of d15 monolayers compared to the control (2.95 Hz, Fig. 3G, Video), confirming that ZNF436 is required to maintain proper cardiomyocyte function.

**Fig. 3.**
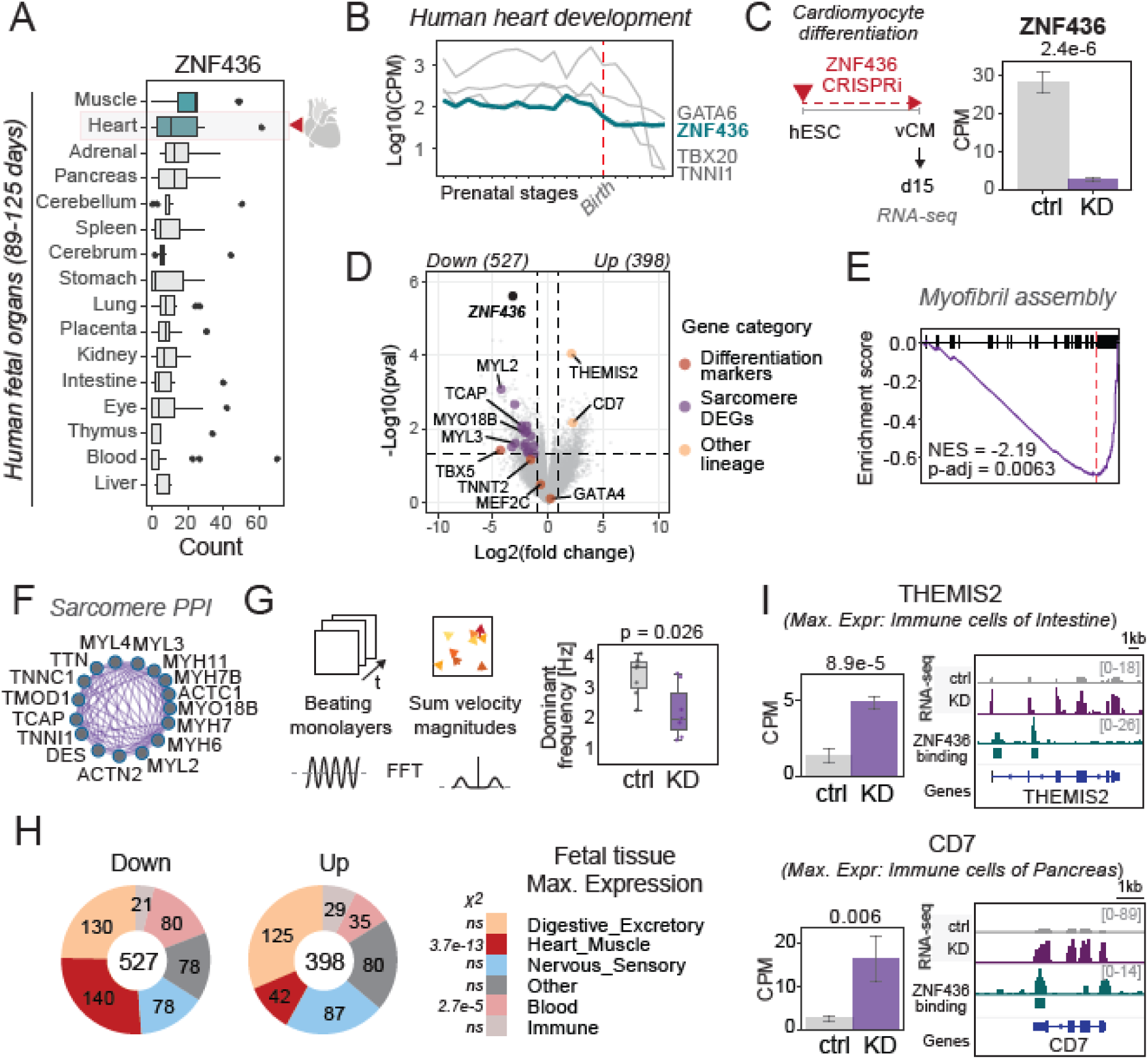
ZNF436 is needed for proper cardiomyocyte contractility. (**A**) *ZNF436* expression in human fetal organs corresponding to 89-125 days of fetal development from Cao, et al. (2020). (**B**) *ZNF436* expression during prenatal and postnatal human heart development stages, together with the expression of *GATA6*, *TBX20* and *TNNI1* from Cardoso-Moreira, et al. (2019). The dashed red line represents the time of birth. CPM, count per million. (**C**) Schematic of ZNF436 KD using CRISPRi during differentiation of human embryonic stem cells (hESCs) to ventricular-like cardiomyocytes (vCMs, d15) with *ZNF436* expression in bulk RNA-seq at d15 cardiomyocytes in the control (grey) and ZNF436 KD (purple) conditions. P-value was computed using a moderated t-test from the limma package. Error bars represent the standard error of the mean (SEM). (**D**) Volcano plot of differentially expressed genes (DEGs) at d15, upon ZNF436 KD. Dashed lines represent significance thresholds for differentially expressed genes (p-value < 0.05 and fold change > 2, n = 3 per condition per stage). P-values were computed using a moderated t-test from the limma package.(**E**) Normalized running enrichment score (NES) for myofibril assembly, derived from Gene Set Enrichment Analysis (GSEA). P-values were computed using permutation testing and adjusted for multiple testing using the Benjamini–Hochberg method to control the false discovery rate (FDR). Terms with an FDR < 0.05 were considered significantly enriched. (**F**) Protein-protein interaction (PPI) network for some of the genes of myofibril assembly downregulated at d15 in ZNF436 KD. (**G**) Schematic of particle image velocimetry (PIV) approach to compute dominant frequencies of d15 vCMs beating monolayers (left). Boxplot showing the dominant beating frequency obtained from Fast Fourier Transform (FFT) analysis of velocity magnitude time series for control and ZNF436 KD condition (right). (n = 7) P-value was computed using the Wilcoxon test. (**H**) Proportions of downregulated and upregulated genes in ZNF436 KD based on the fetal tissue of origin of their maximum expression from Cao, et al. (2020). P-values were computed using the Chi-square test. *X^2^*, chi-square (**I**) Example of ZNF436 binding in the vicinity of upregulated genes, such as *THEMIS2* (top) and *CD7* (bottom), showing bulk RNA-seq coverage upon ZNF436 KD. Bar plots indicate the corresponding expression levels (CPM) for the two genes. (n = 3/condition) P-values were computed using a moderated t-test from the limma package. Error bars represent the SEM.

To investigate whether these effects were mediated by direct ZNF436 targets, we analyzed the genomic proximity of misregulated genes to ZNF436 binding sites (fig. S3F). Unexpectedly, both upregulated and downregulated genes were found near ZNF436 binding sites, supporting a model in which ZNF436 both sustains cardiomyocyte-specific gene expression, and suppresses the activation of alternative transcriptional programs. To test this, misregulated genes were assigned a cell type and a tissue category based on their maximal expression in the human fetal embryo (Cao, et al., 2020). Consistent with our previous analysis, genes downregulated in ZNF436-depleted cells were predominantly expressed in cardiomyocytes and endothelial cells of the fetal heart and muscle (Fig. 3H, and fig. S3G). In contrast, most genes upregulated in this setting showed maximal expression across a wide range of fetal tissues and cell types, with no clear enrichment for a particular set of tissues or cell types of origin (Fig. 3H, and fig. S3G). For example, ZNF436 binding was detected near *THEMIS2* and *CD7*, two genes highly expressed in immune cells of the intestine and pancreas, respectively (Fig. 3I). This indicates that ZNF436 depletion triggers ectopic activation of genes not typically expressed in fetal cardiac cell types. Altogether, these data suggest that ZNF436 safeguards cardiomyocyte identity by maintaining the expression of cardiac-specific genes, while preventing the activation of transcriptional programs from diverse non-cardiac origins.

### ZNF436 restricts chromatin accessibility in cardiomyocytes

Since ZNF436 binding was detected near both upregulated and downregulated genes, we investigated the underlying molecular mechanisms driving this phenotype. Given its high sequence conservation, we hypothesized that the vKRAB domain of ZNF436 was the functional module responsible for the recruitment of additional chromatin effectors (Fig. 2B). To test this, we employed a proximity biotinylation assay in which the biotin ligase TurboID was fused to either a full-length ZNF436 (ZNF436-TID) or its KRAB-deleted version (ΔKRAB-TID), alongside a nucleus-targeted GFP control (GFP-TID) (Fig. 4A, and fig. S4A) (Santos-Barriopedro, et al. 2023). Stable hESC lines expressing these constructs were differentiated into vCMs, followed by biotin pull-down and mass spectrometry at d15 (Fig. 4A). Both ZNF436-TID and ΔKRAB-TID exhibited predominant nuclear localization, followed by the enrichment of predominantly chromatin-associated proteins in their vicinity (Fig. 4, B and C, and fig. S4B). Although similar proteins were detected near both constructs, the vKRAB domain mediated the enrichment of 66 proteins that were absent in its absence (Fig. 4B, and fig. S4, C and D), underscoring its role as a key functional effector module. Among these vKRAB-dependent ZNF436 neighbors, the members of the TRIM family, TRIM24 and TRIM33, stood out (Fig. 4C). Both proteins are related to TRIM28, but function as chromatin readers operating independently of H3K9me3 (Appikonda, et al. 2018; Sekirnik, et al. 2022). In contrast, TRIM28 was not associated with ZNF436 (TRIMs, Fig. 4C), as verified by co-immunoprecipitation in A673 sarcoma cell line (fig. S4E). These findings suggest that ZNF436 vKRAB might interact with non-canonical TRIM proteins, supporting a regulatory mechanism distinct from TRIM28-mediated heterochromatin formation. In line with a role in active gene regulation, vKRAB-enriched proteins included other TFs, subunits of the RNA polymerase II, and mRNA-associated proteins (Fig. 4C, and fig. S4F). Most notably though, the largest group of proteins belonged to the components of ATP-dependent chromatin-remodeling complexes (n = 16), with an overrepresentation of members from SWI/SNF complexes (Fig. 4C). Among these, we identified subunits specific to the canonical BAF complex, such as ARID1A and DPF2, and the “polybromo-associated” BAF complex (PBAF), such as PBRM1 and BRD7 (Baxter, et al. 2023). Importantly, ΔKRAB-TID showed a reduced association to both PBRM1 and BRD7, suggesting that the vKRAB steers the composition of the SWI/SNF complex toward PBAF (Fig. 4D).

**Fig. 4.**
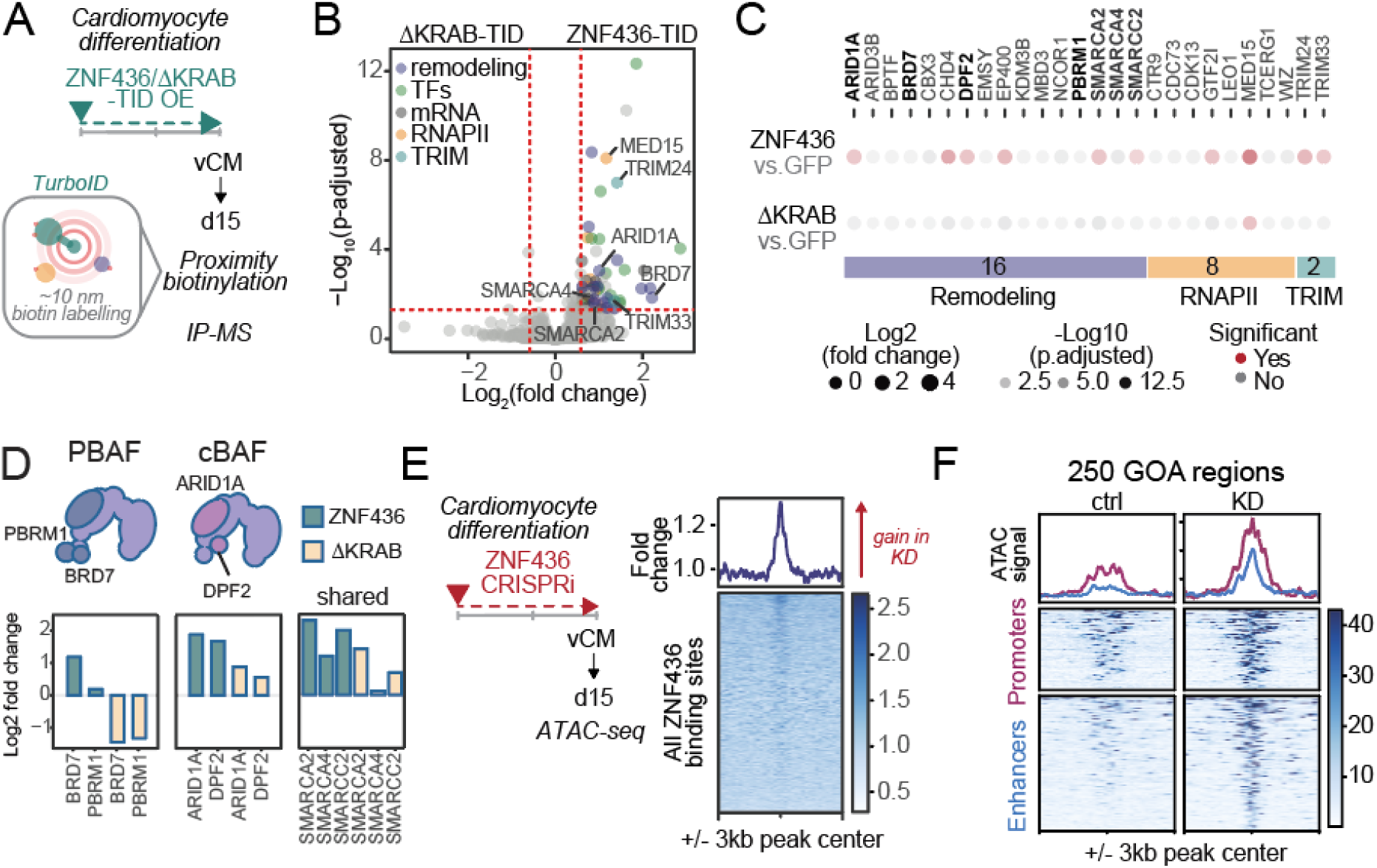
ZNF436 restricts accessibility in cardiomyocytes. (**A**) Schematic representation of the proximity labeling experiment for identifying proteins enriched in the ZNF436 vicinity in d15 cardiomyocytes. TID, turboID. (**B**) Volcano plot of differentially enriched proteins between ZNF436-TID and ΔKRAB-TID in d15 cardiomyocytes. Dashed lines represent significance thresholds (p-adjusted < 0.05 and fold change > 1.5, n = 6 replicates). P values were computed using a moderated t-test from the limma package, corrected for multiple testing using the Benjamini-Hochberg’s method. (**C**) Dotplot showcasing individual proteins enriched for ZNF436-TID highlighted in Fig.4B. Fold change, significance and p-adjusted values were derived from comparisons of the indicated condition with the GFP-TID control at d15 (p-adjusted < 0.05). P values were computed using a moderated t-test from the limma package, corrected for multiple testing using the Benjamini-Hochberg’s method. (**D**) Schematic representation of SWI/SNF cBAF and PBAF complexes (top), and the log2 (fold change) detected for ZNF436-TID or ΔKRAB-TID *vs.* GFP-TID for the indicated subunits (bottom). (**E**) Schematic representation for profiling chromatin accessibility using ATAC-seq upon ZNF436 KD (left) and a heatmap representing the fold change of ATAC-seq accessibility (ZNF436 KD over control condition) centered on all ZNF436 peaks (n=3633 regions, right). (**F**) Heatmap of accessibility based on ATAC-seq data for 250 high-confidence ZNF436-bound regions with significant gain of accessibility (GOA), in control and ZNF436 KD condition, split into promoter and enhancer regions. P-values were computed using the t-test.

As the cell-type specific composition of the BAF complex was shown to regulate stage-specific chromatin accessibility during cardiomyocyte differentiation (Alexander, et al. 2015; Hota et al., 2019; Hota, et al. 2022), we performed ATAC-seq in d15 vCMs upon ZNF436 KD (Fig. 4E) (Buenrostro, et al., 2013). Despite bidirectional changes in gene expression, ZNF436 binding sites showed a consistent and widespread increase in chromatin accessibility in ZNF436-depleted vCMs versus control (fold change >1, Fig. 4E). Following this general trend, 95% (250/264) of significantly differentially accessible regions overlapping ZNF436 peaks gained accessibility upon ZNF436 KD (GOA regions, fig. S4G). Notably, GOA regions were characterized by a reduced accessibility in the control condition (Fig. 4F), consistent with a model in which ZNF436 co-occupies these loci with PBAF complexes at sites lacking active remodeling, similar to REST (Grossi, et al., 2025). This pattern was observed in GOA associated with promoters, as well as enhancers, including those located in promoter-distal genic and intergenic areas (Fig. 4F, and fig. S4G). Together, these findings reveal that ZNF436 restricts chromatin accessibility at its binding sites, likely through selective engagement with specific SWI/SNF complexes and alternative TRIM proteins.

### Ectopic regulatory activity of L2/MIRs is limited by ZNF436

We next examined the functional consequences of increased chromatin accessibility at GOA regions by profiling the epigenomic landscape of ZNF436-depleted vCMs using Cut&Tag (Kaya-Okur, et al., 2019). As expected, no change in H3K9me3 levels was detected at these sites, confirming that ZNF436 functions independently of heterochromatin deposition (fig. S5A). On the other hand, significant alterations were observed for H3K27ac, H3K4me1 and H3K4me3 (Fig. 5A, and fig. S5B). These changes were consistent with the underlying genomic features, as enhancers showed increased H3K27ac and H3K4me1, while promoters gained both H3K27ac and H3K4me3 (Fig. 5A). Together, these results indicate that ZNF436 binding restricts ectopic *cis*-regulatory activity of L2/MIRs in cardiomyocytes.

**Fig. 5.**
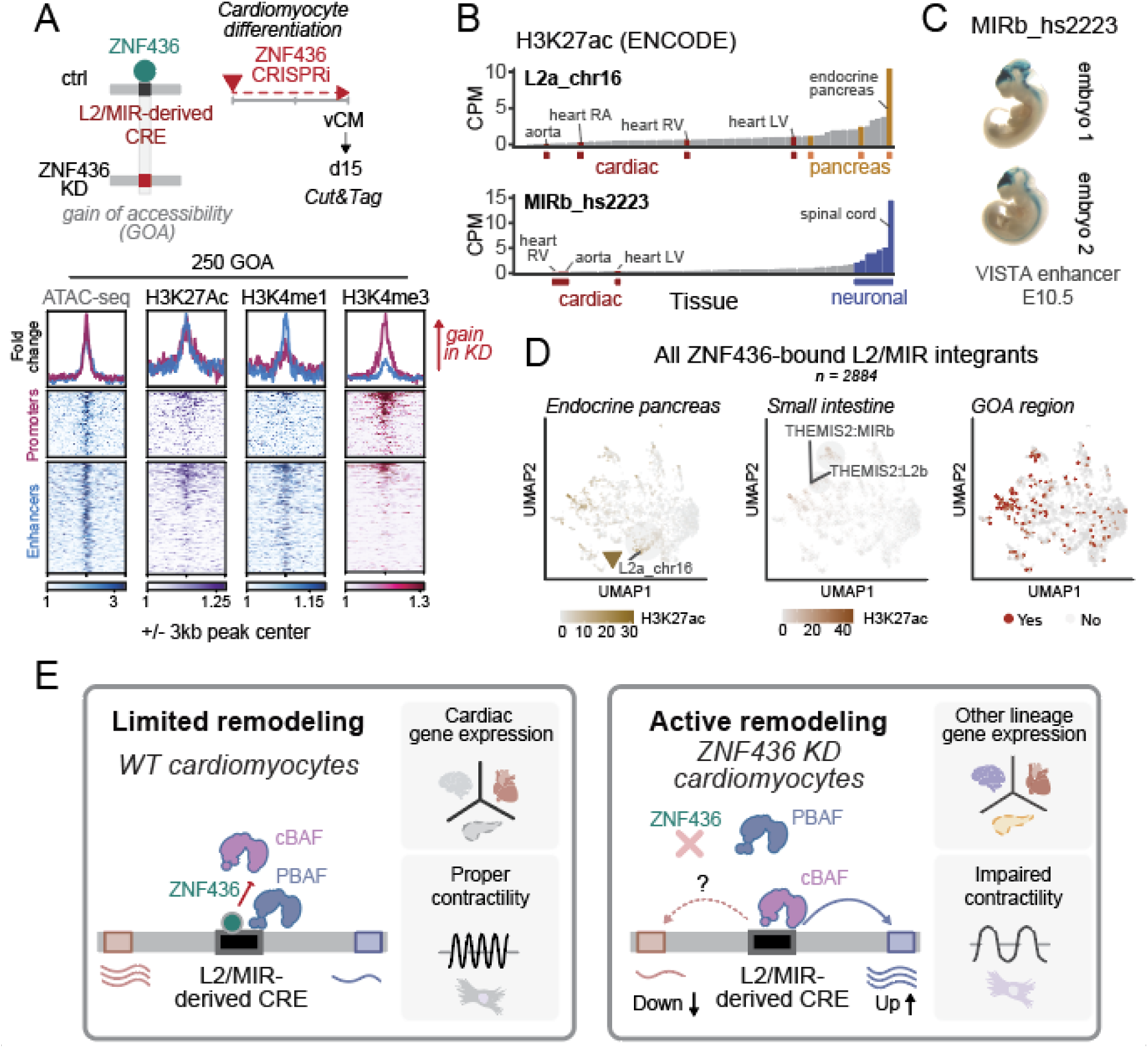
ZNF436 restricts L2/MIR-derived CRE activity in cardiomyocytes. (A) Schematic representation for profiling promoter- and enhancer-associated histone marks upon ZNF436 KD in d15 vCMs (left). with heatmaps representing the fold change for ATAC-seq signal for GOA regions, alongside the fold change of Cut&Tag signal for H3K27ac, H3K4me1 and H3K4me3 (ZNF436 KD over control condition) centered on GOA regions, split into promoter and enhancer regions. (**B**) H3K27ac signal across 67 tissues from ENCODE for the ZNF436-bound L2a_chr16 and MIRb_hs2223 integrants. CPM, count per million. (**C**) Representative E10.5 transgenic embryos showing whole-mount LacZ staining, with LacZ reporter expression (blue) driven by the MIRb-derived hs2223 element (Visel, et al. 2007). (**D**) UMAP of all L2/MIR integrants overlapping ZNF436 peaks (n=2884) based on the ENCODE H3K27ac signal across tissues with a projection of H3K27ac signal from endocrine pancreas and hippocampal brain region, and annotation of specific L2/MIR integrants overlapping GOA ZNF436 peaks (red). (**E**) Model depicting the physiological role of L2/MIR-derived CREs, whose activity is restricted by ZNF436 in cardiomyocytes through SWI/SNF remodeling. ZNF436 depletion in cardiomyocytes results in ectopic CRE activity leading to misregulated genes affecting cardiomyocyte contractility.

To delineate the activity of these CRE in a broader physiological context, we derived a maximal CRE activity score of ZNF436-bound TE integrants, based on their H3K27ac levels across 67 tissues, using Z-score normalization (See methods, fig. S5C). This relative specificity score allowed to identify GOA L2/MIR integrants endowed with a high tissue-specific activity (specificity score > 0.67, fig. S5C). Among those, L2a_chr16 showed strong tissue-specific enhancer activity in the endocrine pancreas (Fig. 5B, specificity score = 0.77). The ectopic activation of this enhancer upon ZNF436 KD in vCMs mimicked its active state in the pancreas, but not in the heart or brain (fig. S5D). Similar results were observed for the neural-specific MIRb_hs2223 (specificity of 0.82 in spinal cord, Fig. 5B and fig. S5D), the activity of which was documented *in vivo* at E10.5 of mouse development (VISTA enhancer, Fig. 5C) (Visel, et al., 2007).

Building on these observations, we extended the analysis to all ZNF436-bound L2/MIR elements based on their H3K27ac profiles across tissues. Expanding on the previously defined KZFP-driven UMAP structure of TE subfamilies (Fig. 1B), we found that similarities in their tissue-specific CRE activity shaped the spatial organization of L2/MIR integrants. This revealed distinct sets of tissue-specific activities. For example, a pancreas-associated group containing L2a_chr16, as well as groups associated with intestine, muscle, or brain, suggested that ZNF436-bound L2/MIR integrants function as active CREs in diverse tissues (Fig. 5D and fig. S5, E and F). Consistently, high-confidence GOA regions were dispersed across the UMAP space, indicating no preferential association with any specific tissue-defined group (Fig. 5D). By projecting the fold change in H3K27ac signal in ZNF436-depleted vCMs versus controls, we observed widespread ectopic CRE activation, and on selected instances, we could link CRE activation with the ectopic expression of non-cardiac genes identified in RNA-seq analysis (Fig. 5D, and fig. S5, F-H). For example, *THEMIS2*, previously identified as highly expressed in fetal intestine and upregulated upon ZNF436 KD, was linked to MIRb and L2b integrants located in the region of high H3K27ac signal in the small intestine (Fig. 3I and 5D, and fig. S5G). Similar results were observed for *PACSIN1*, with nearest L2c showing high activity in the angular gyrus brain region (fig. S5F). Accordingly, ZNF436 emerges as a critical regulator of cardiac identity, maintaining the expression of cardiac gene programs by constraining CRE activity linked to alternative tissue lineages.

## DISCUSSION

KZFPs are well-known repressors of TE-derived regulatory activity through TRIM28-mediated H3K9me3 deposition. However, whether KZFPs operate beyond this canonical pathway to modulate CREs remains largely unexplored. In this study, we identify ZNF436 as a KRAB-containing KZFP that regulates L2/MIR-derived CREs *via* a noncanonical mechanism, one that relies on the restriction of chromatin accessibility through interactions with the SWI/SNF complex, offering a more dynamic and context-specific form of regulatory control.

We provide a comprehensive map of epigenomic states across KZFP-bound TEs, with a focus on the mammalian-specific L2/MIR elements. Although these TEs have been previously associated with enhancer and promoter activities (Jjingo et al., 2014; Cao et al., 2019; Roller et al., 2021), our work uncovers new layers of regulatory stratification shaped by KZFP binding, chromatin context, and tissue-specific activity. With around half a million integrants in the human genome, L2/MIRs do not function as a uniform regulatory unit. Instead, we show that their CRE activity can be broadly delineated through distinct KZFP binding profiles. Notably, despite being bound by numerous KZFPs, L2/MIRs exhibit relatively low H3K9me3 enrichment (Fig. 1D). This pattern is associated with their interaction with a subset of KZFPs carrying vKRAB domains, which do not recruit TRIM28 and whose functions remain largely unexplored (Tycko, et al., 2023; Rosspopoff and Trono, 2023). Among these, we identify ZNF436 as a highly conserved vKRAB-containing KZFP that emerged alongside the expansion of L2/MIR elements ∼160 million years ago and exhibits exceptional binding specificity toward these elements across mammalian species (Fig. 2). ZNF436 is highly expressed in the developing fetal heart *in vivo*, and its depletion in hESC-derived cardiomyocytes leads to widespread gene expression changes and impaired cardiomyocyte function (Fig. 3).

Mechanistically, ZNF436 plays a dual role in maintaining cardiomyocyte function, where it sustains the expression of cardiomyocyte-specific genes while preventing the activation of genes associated with alternative lineages. This ectopic gene activation is mirrored at the chromatin level, where ZNF436 limits the accessibility of L2/MIR-derived sequences and attenuates their CRE activity. Our work provides direct evidence for a vKRAB-dependent mechanism that restricts CRE accessibility through selective interactions with the SWI/SNF chromatin remodeling machinery, thereby marking a significant extension of the classical KZFP paradigm. In cardiomyocytes, the vKRAB may facilitate ZNF436 co-occupancy with the PBAF subtype of SWI/SNF complexes (Fig. 4). This is consistent with recent findings that PBAF contributes to the low accessibility within inactive chromatin environments to enable the binding of repressive TFs, such as REST (Grossi, et al. 2025). In turn, loss of REST has been linked to increased chromatin accessibility, ectopic CRE activation, and aberrant neuronal gene expression (Soleimani, et al. 2024). Similarly, ZNF436 depletion results in a marked gain in chromatin accessibility, along with the acquisition of histone modifications characteristic of active enhancers and promoters (Fig. 5A). Strikingly, the CREs activated upon ZNF436 loss resemble those active in a variety of tissues, including brain and pancreas, suggesting that ZNF436 does not repress a single alternative lineage, but instead prevents multilineage CRE activation. In line with this, upregulated genes upon ZNF436 KD showed no enrichment for a specific tissue origin (Fig. 3). Together, these findings support a model in which ZNF436 partakes in defining SWI/SNF subunit composition at poised and repressed TE-derived CREs, thereby safeguarding cardiomyocyte identity and function. Importantly, the gradual induction of ZNF436 during cardiomyocyte differentiation, suggests that it coordinates the chromatin closure of actively remodeled sites into a low-accessibility configuration through selective engagement with the PBAF complex. It remains to be addressed whether PBAF follows a temporally regulated chromatin association pattern during cardiomyocyte differentiation, similar to canonical BAF complexes (Alexander, et al. 2015, Hota, et al. 2019, Hota, et al. 2022) and to which extent ZNF436 influences this process. In addition to SWI/SNF complexes, our data show that ZNF436 interacts in a vKRAB-dependent manner with alternative TRIM proteins, including TRIM24 and TRIM33. TRIM24, in particular, has been shown to localize to closed or low-accessibility chromatin regions in a TF-dependent manner, which facilitates rapid chromatin opening under stress conditions (Isbel, et al., 2023). Therefore, the association of ZNF436 with TRIM24 may reflect a more flexible, primed regulatory state of CREs that remains responsive to developmental signals. We propose that, unlike a rigid H3K9me3-based repression, ZNF436-mediated recruitment of TRIM24 and/or the PBAF complex enables a more dynamic and reversible layer of CRE regulation, that primes chromatin for downstream cellular or signaling cues.

Nevertheless, several questions remain. We have yet to broadly link the ectopic activation of CREs to specific misregulated genes, and further explore the implication of ZNF436 binding in the vicinity of cardiomyocyte-specific genes. For instance, the presence of the components of RNA polymerase II and adjacent proteins in the vicinity of ZNF436 in our proximity labelling experiment could suggest more than one mechanism for ZNF436-mediated regulation (Fig. 4). This would suggest not only context-specific functions for ZNF436, but also locus-specific regulatory grammar. Further investigation, including capture techniques and locus-specific proteomics, will be required to better understand how specific CREs influence gene expression in a context-dependent manner. Moreover, our current analysis is limited to day 15 of cardiomyocyte differentiation, when ZNF436 expression peaks, leaving open questions about the temporal onset and progression of its chromatin regulatory activity and how it precedes or accompanies gene expression changes.

While our work focuses on ZNF436, it raises broader questions about the functional diversity of vKRAB domains. Although recent studies suggest that other vKRAB-containing KZFPs also act independently of H3K9me3 (Moore et al., 2025; Rosspopoff et al., 2025), the presence of accessory domains such as DUF3669 and SCAN may contribute additional functional specificity (Begnis et al., 2024; Matsushima et al., 2025). Importantly, vKRAB sequences vary substantially across human KZFPs (Helleboid, et al., 2019; Tycko, et al. 2020), and many appear to be degenerating (Emerson and Thomas, 2011), suggesting functional divergence within the vKRAB family.

In sum, our data suggest that over long evolutionary time periods, the relationship between KZFPs and TEs evolves to enable a more reversible, dynamic mechanism of CRE restriction. Our findings highlight the importance of dynamically regulated CRE activity in balancing the repression of alternative lineages with the activation of lineage-specific programs. This flexibility in chromatin state regulation supports regulatory plasticity during development and differentiation, reinforcing the idea that cell fate is not fixed but follows a dynamic and responsive trajectory. Disruption of this balance may impair the proper acquisition or maintenance of cell identity. Extending this concept, together with others (Plaisance, et al., 2023), our work reveals that ancient, mammalian-specific TEs actively contribute to mammalian heart development. These TE-derived enhancers have been integrated into gene regulatory networks that orchestrate cardiac differentiation and function, potentially adding a layer of regulatory complexity and evolutionary adaptability to the mammalian cardiovascular system.

## Contributions

D.M. conceptualized the project, performed the experiments, analyzed the data, and wrote the manuscript. O.R. and D. T. conceptualized the project and wrote the manuscript. D.T. secured the funding. J.D. analyzed the data and wrote the manuscript. W.M. performed the ATAC-seq experiment. R. H. processed the proteomic samples and wrote the manuscript. E.P. helped with genomic data processing. S.O. helped with the cloning, cell culture, and lentivector production. All authors approved the final manuscript.

## Acknowledgments

We thank Martina Begnis, Olimpia Bompadre, Iris Arianna Dorschel and Cedric Feschotte for reading the manuscript, fruitful discussions and feedback. We specially thank Romain Forey for his insightful feedback during manuscript preparation. We thank members of Gene Expression Core Facility (GECF), Proteomics Core Facility (PCF) and Bioimaging and Optics Platform (BIOP) of EPFL for their help, suggestions and expertise in sample processing. We thank Enzo Guialardon and other past member of the lab for their help.

## Funding

This work was funded by the European Research Council (KRABnKAP no. 268721; Transpos-X no. 694658), the Swiss National Science Foundation (grants 310030_192613 and 310030_188803), and the Aclon Foundation, all awarded to DT.

## Conflict of interests

The authors declare no competing interests.

---

**Figure S1.**
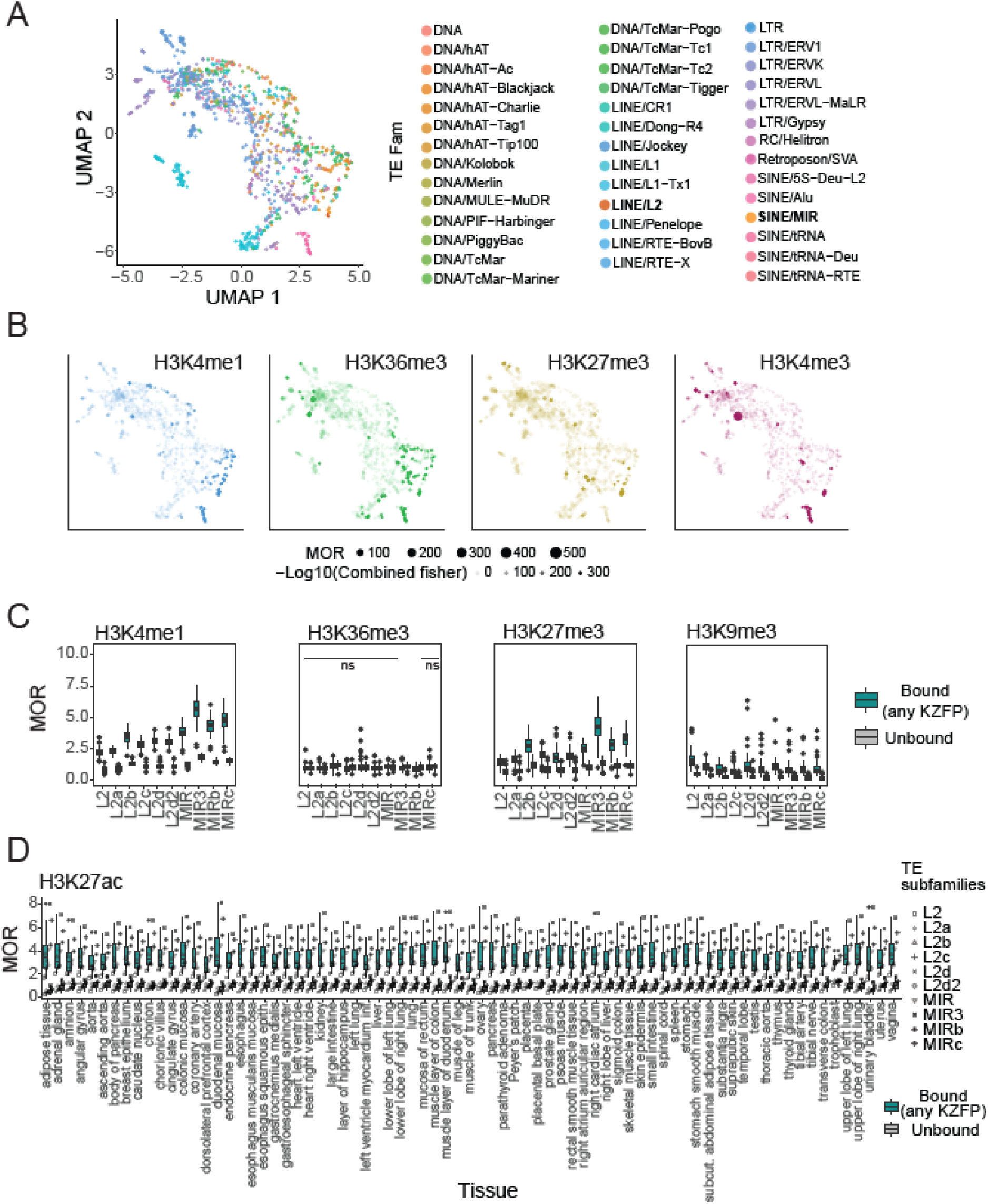
Chromatin states define the distinct behavior of KZFP-bound TE subfamilies. (**A**) UMAP representation of Giggle scores for each TE subfamily, colored by TE family. (**B**) UMAP projection of the MOR enrichment score for indicated histone marks. (**C**) MOR for indicated histone marks for L2/MIR subfamilies across tissues, grouped by KZFP-bound and unbound integrants. All KZFP-bound integrants were significantly enriched compared to unbound ones (p-value < 0.001) unless highlighted by ns (not significant). P-values computed using a t-test. (**D**) MOR for H3K27Ac for L2/MIR subfamilies per tissue, grouped by KZFP-bound and unbound integrants. All KZFP-bound integrants were significantly enriched compared to unbound ones (p-value < 0.001). P-values were computed using a paired t-test.

**Figure S2.**
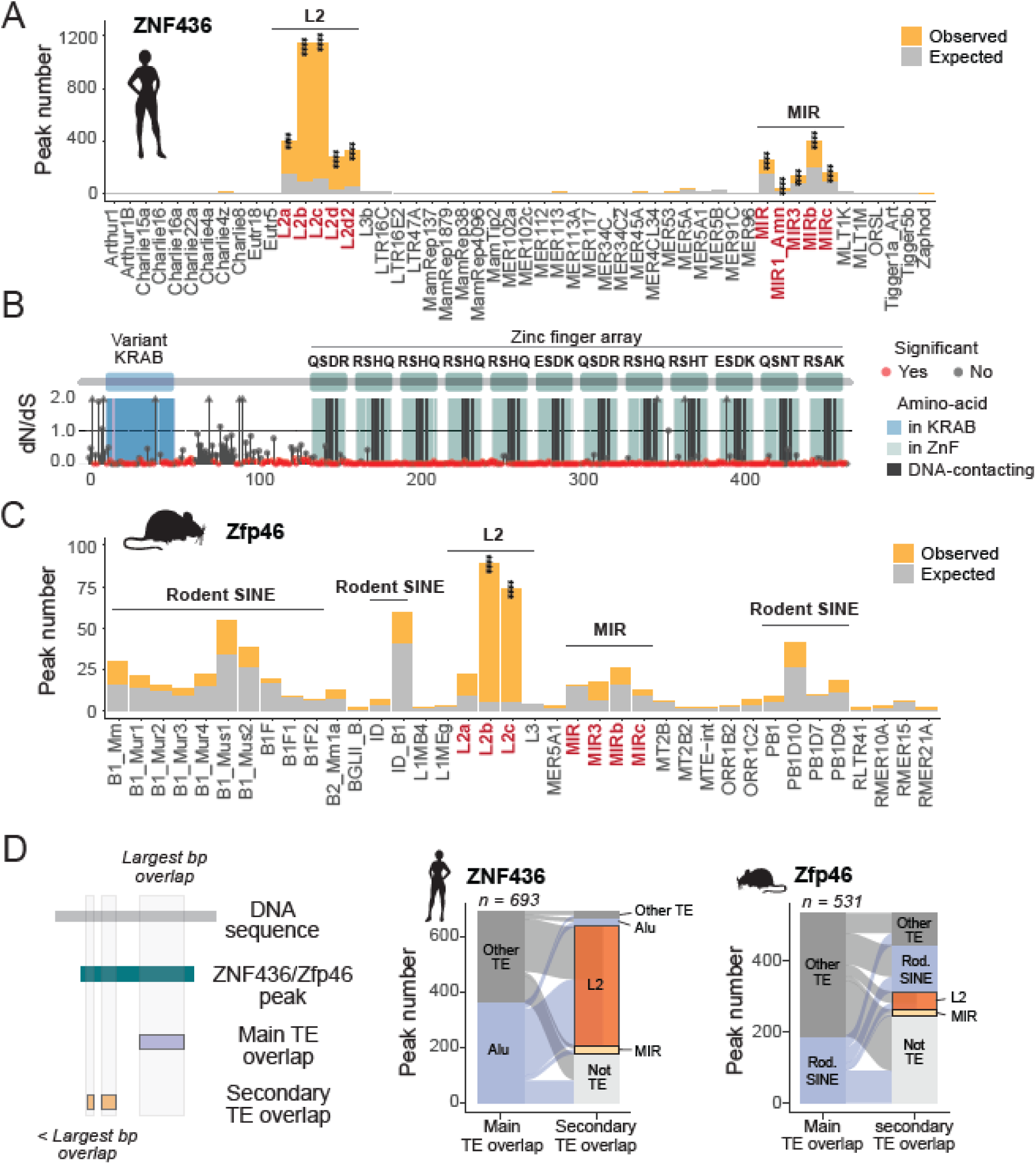
Both human ZNF436 and mouse Zfp46 orthologs are enriched for L2/MIR binding. (**A**) Enrichment of TE subfamilies overlapping human ZNF436 peaks. P-values were computed using a binomial test, adjusted for multiple testing using the Benjamini-Hochberg method. (**B**) Schematic representation of the dN/dS ratio computed for each amino acid of ZNF436 over the indicated list of domains. Red dots indicate residues with a significant p-value below 0.05. P-values were computed using a Mann-Whitney U test. (**C**) Enrichment of TE subfamilies overlapping mouse Zfp46 peaks. Rod. SINE, rodent-specific SINE elements. P-values were computed using a binomial test, adjusted for multiple testing using the Benjamini-Hochberg method. (**D**) Schematic outlining the strategy used to assign main and secondary TE subfamilies based on the extent of base-pair overlap with ZNF436/Zfp46 peaks (left). Alluvial plot showing secondary TE overlaps for ZNF436/Zfp46 peaks whose main overlap was with TE subfamilies other than L2/MIR (right).

**Figure S3.**
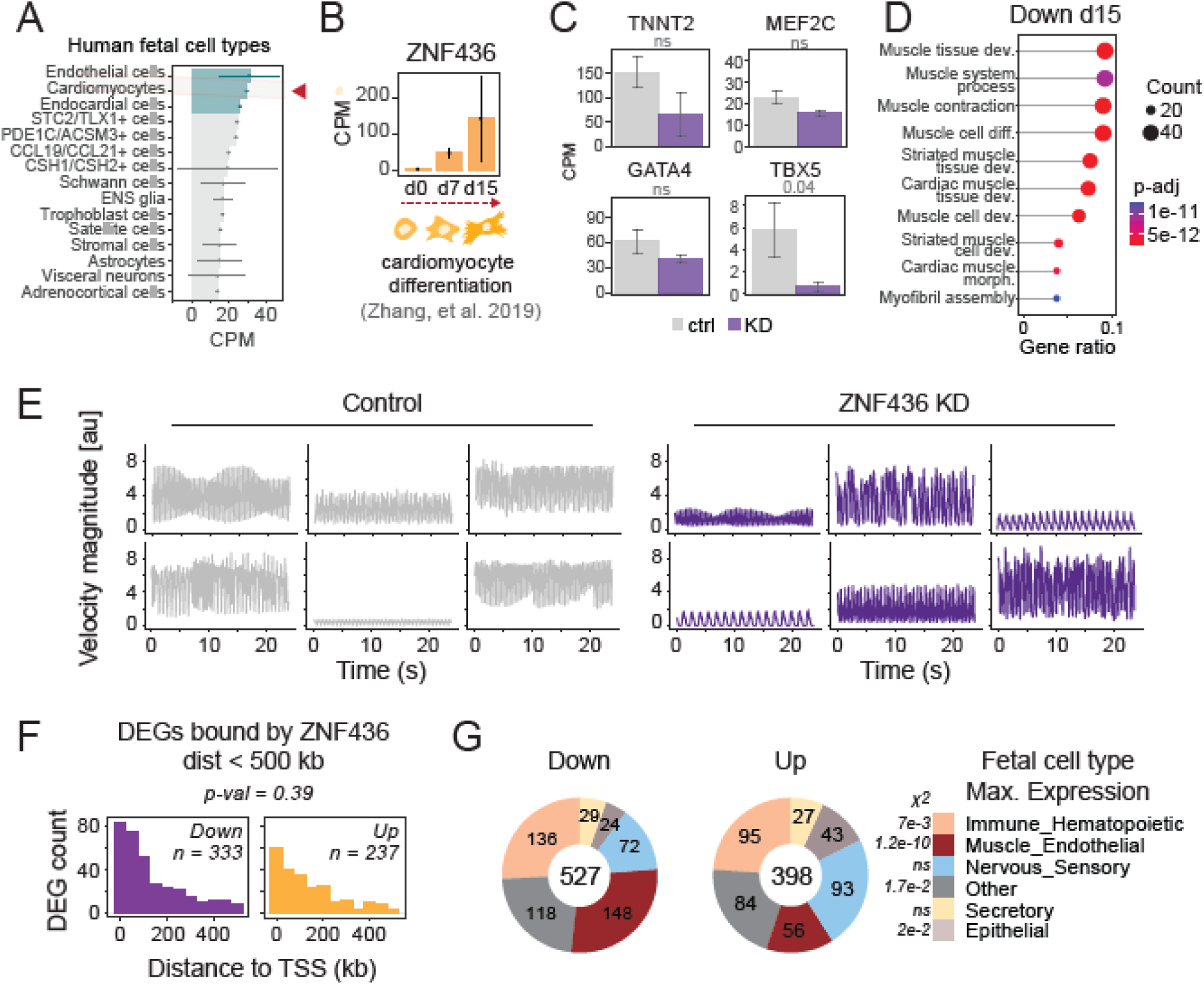
ZNF436 KD affects cardiomyocyte functionality. (**A**) *ZNF436* expression in human fetal cell types from Cao, et al. (2020), corresponding to 89-125 days of fetal development. (**B**) *ZNF436* expression during cardiomyocyte differentiation from Zhang, et al. (2019). (**C**) Expression of known vCM markers in control vs. ZNF436 KD cardiomyocytes. (n = 3/condition) P-values were computed using a moderated t-test from the limma package. Error bars represent the SEM. (**D**) Enriched ontologies for genes significantly downregulated in ZNF436 KD. P-values were computed using a hypergeometric test, and adjusted for multiple testing using the Benjamini– Hochberg procedure to control the FDR. GO terms with an FDR < 0.05 were considered significantly enriched. (**E**) Individual velocity magnitude time series on which Fast Fourier transform (FFT) was performed for obtaining dominant frequencies for control and ZNF436 KD conditions. (**F**) Distances to the nearest ZNF436 peaks for both downregulated and upregulated genes upon ZNF436 KD, limited to a 500 kb window as described in Pulver, et al. (2023). P-value was computed using the Wilcoxon test. (**G**) Proportions of downregulated and upregulated genes in ZNF436 KD based on the fetal cell type of origin of their maximum expression from Cao, et al. (2020). P-values were computed using the Chi-square test. *X^2^*, chi-square

**Figure S4.**
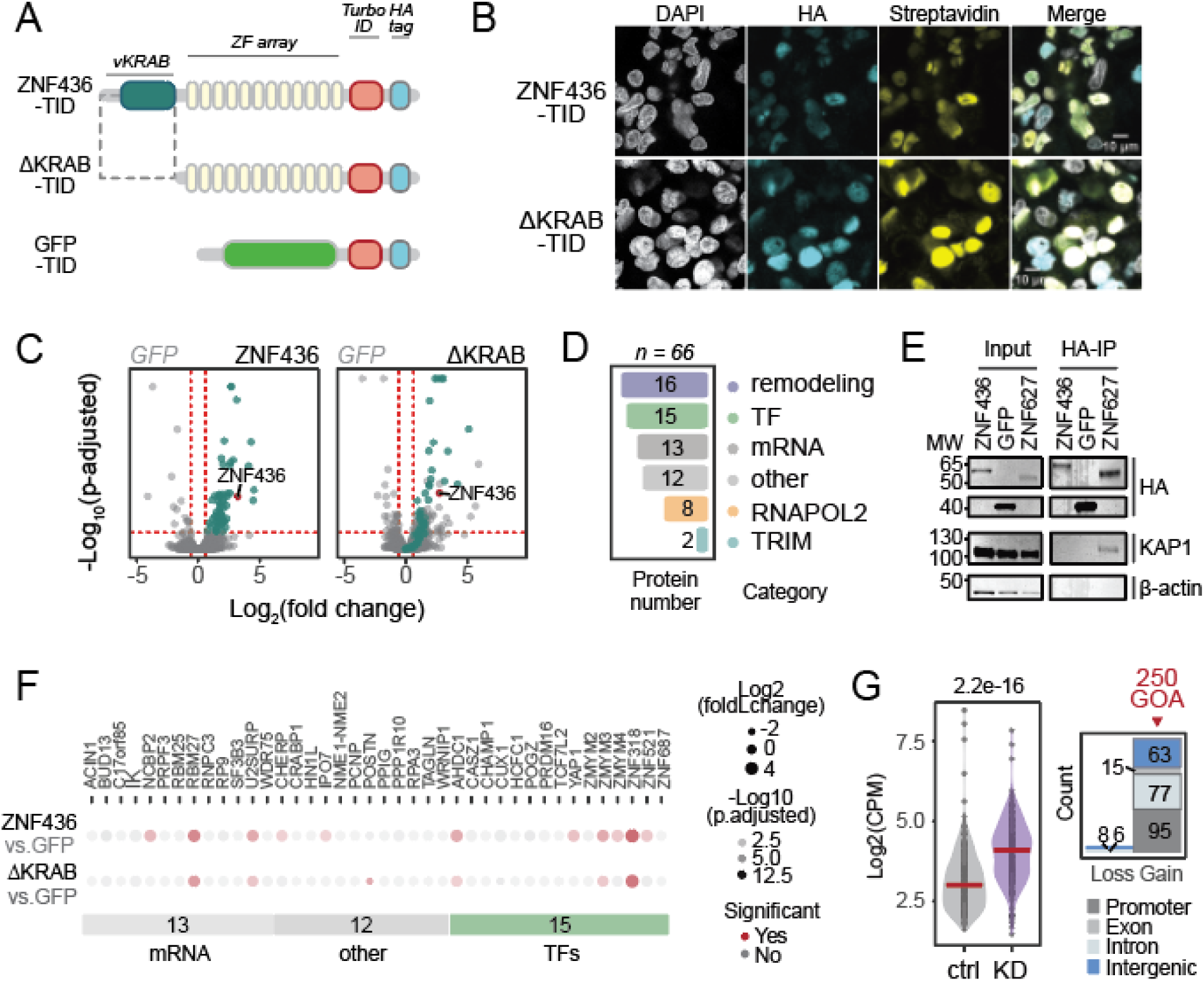
ZNF436 proximal proteins are restricted to cardiomyocytes. (**A**) Schematic representation of HA-tagged ZNF436-TID, ΔKRAB-TID, and GFP-TID constructs used for stable overexpression in hESC-derived ventricular cardiomyocytes (vCMs) for proximity labeling experiments. TID, TurboID. (**B**) Immunofluorescence experiments showing nuclear localization of HA-tagged ZNF436-TID and ΔKRAB-TID at d15, as well as the overlap with streptavidin signal indicating nuclear biotinylation (scale bar = 10 μm). (**C**) Volcano plots of protein enrichment for ZNF436-TID and ΔKRAB-TID *vs.* GFP-TID. The cyan-colored dots represent proteins enriched in the ZNF436-TID *vs.* GFP-TID condition. Dashed lines represent significance thresholds (p-adjusted < 0.05 and fold change > 1.5, n = 3). P values were computed using a moderated t-test from the limma package, corrected for multiple testing using the Benjamini-Hochberg’s method. (**D**) Bar plot showing the categories assigned to the 66 enriched proteins in ZNF436-TID vs. ΔKRAB-TID at d15. (**E**) Co-immunoprecipitation of HA-tagged constructs showing the lack of interaction of vKRAB-containing ZNF436 with KAP1, contrary to the standard KRAB-containing KZFP, ZNF627, performed in the A673 cell line. (**F**) Dotplot showcasing individual proteins enriched for ZNF436-TID highlighted in Fig.4B. Fold change, significance and p-adjusted values were derived from comparisons of the indicated condition with the GFP-TID control (p-adjusted < 0.05). P values were computed using a moderated t-test from the limma package, corrected for multiple testing using the Benjamini-Hochberg’s method. (**G**) ATAC-seq signal over ZNF436-bound regions identified as significantly differentially accessible at d15 in ZNF436 KD (n = 264) and genomic features associated with regions with significant gain and loss of accessibility. GOA, Gain Of Accessibility. P-value was computed using a paired t-test.

**Figure S5.**
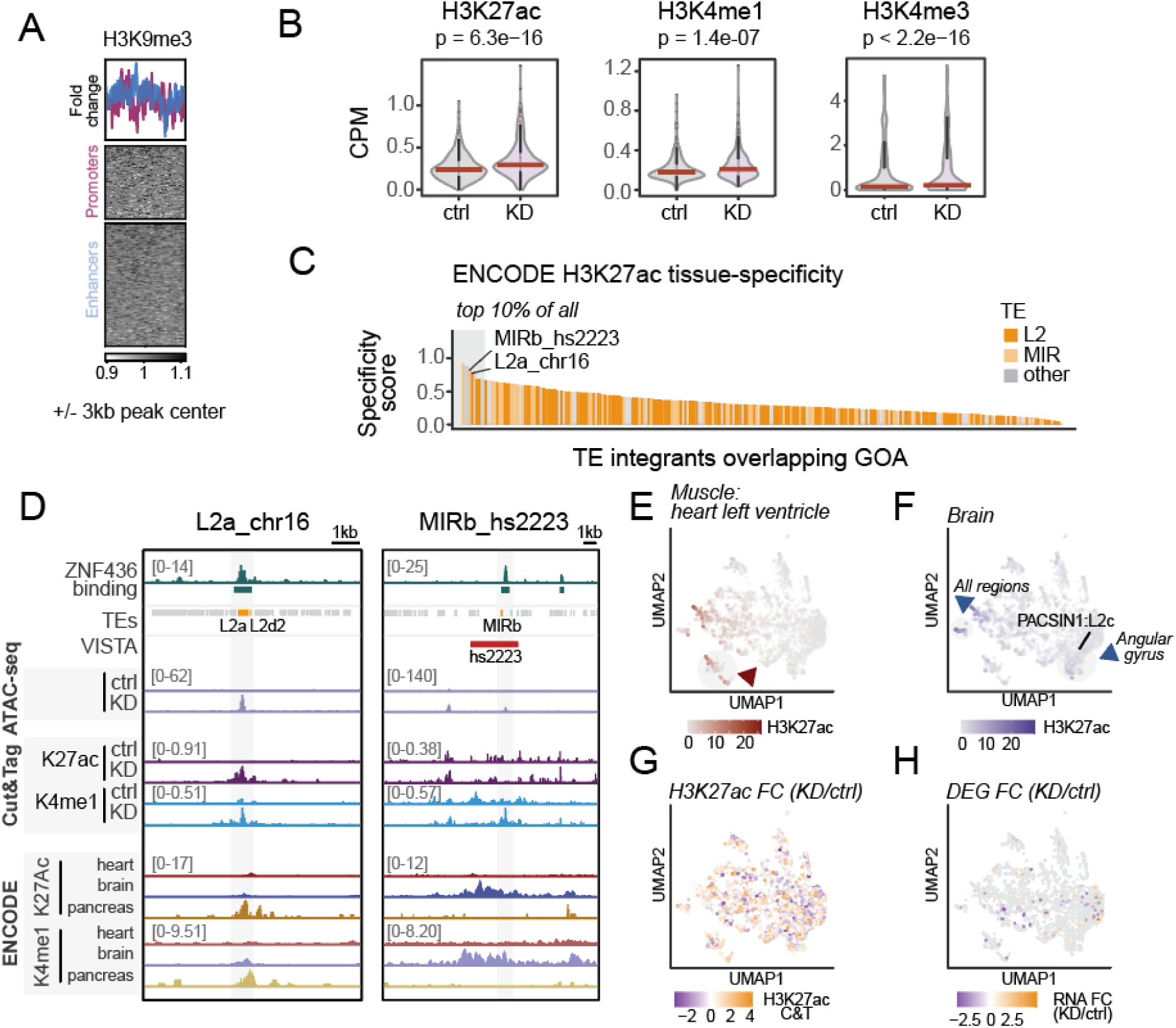
ZNF436 dictates the tissue-specific activity of L2/MIR-derived CREs. (**A**) Heatmap representing the fold change for H3K9me3 Cut&Tag signal fold change (ZNF436 KD over control condition) centered on GOA regions, split into promoter and enhancer regions. (**B**) Quantification of H3K27ac, H3K4me1 and H3K4me3 changes over 250 GOA regions in ZNF436 KD versus control condition at d15 (n = 250). P-values were computed using a paired t-test. (**C**) ENCODE H3K27Ac-based relative tissue-specificity scores for ZNF436-bound TE integrants within GOA regions, with emphasis on the L2a_chr16 and MIRb_hs2223 L2/MIR integrants. Top 10% calling is based on all TE integrants within 3633 peaks. (**D**) Genomic regions containing L2a_chr16 (left) and MIRb_hs2223 (right), with tracks representing signals from ATAC-seq and Cut&Tag upon ZNF436 KD, as well as tissues from ENCODE. (**E-H**) UMAP of all L2/MIR integrants overlapping ZNF436 peaks (n=2884) based on the ENCODE H3K27ac signal across tissues with a projection of H3K27ac signal from heart left ventricle (E), brain region angular gyrus (F), H3K27ac signal fold change over 2884 L2/MIR integrant in ZNF436 KD over control condition (G), and RNA-seq CPM fold change in ZNF436 KD over control for assigned misregulated genes to 2884 L2/MIRs (n = 131).

## Methods

### Cell culture and vCM differentiation

Human embryonic stem cell (hESC) line H9 was grown at 37°C in 95% air, 5% CO2 in mTeSR plus media (STEMCELL Technologies, 100-0276) on plates previously coated with Matrigel (Corning, 354277). Cells were routinely passaged every 4 days using gentle cell dissociation reagent (STEMCELL Technologies, 100-0485) for 5 min at room temperature for clump passaging. To obtain a single-cell suspension, cells were incubated for 10 min at 37°C using gentle cell dissociation reagent and incubated for 24h with mTeSR plus supplemented with 10 μM of ROCK inhibitor (STEMCELL Technologies, 72307). hESCs were differentiated into cardiomyocytes using a ventricular cardiomyocyte differentiation kit (STEMCELL Technologies, 05010) following the manufacturer’s protocol. At day 15 of differentiation, ventricular-like cardiomyocytes (vCMs) were dissociated using 0.25% trypsin (Thermo Fisher Scientific, 25300054) for 5 min at 37°C, which was quenched using cardiomyocyte support media (STEMCELL Technologies, 05027).

HEK293T and A673 sarcoma cell lines were maintained in Dulbecco’s Modified Eagle’s Medium (DMEM, Gibco) supplemented with 10% fetal bovine serum (FBS, Hyclone) and 1% penicillin/streptomycin (Gibco). Cells were routinely passaged every 5 days using 0.25% trypsin (Gibco) for 5 min at 37°C, following a 1:10 dilution.

### Lentivirus production

Lentivectors were generated as previously described in (Begnis, et al., 2024) using HEK293T cells.

### CRISPRi experiments

The CRISPRi system was used to inhibit ZNF436 transcription in primed H9 hESCs. DNA oligonucleotides corresponding to the sgRNAs sequences were designed using CRISPick software (https://portals.broadinstitute.org/gppx/crispick/public), maintained by the Broad Institute. Oligonucleotide pairs were annealed to generate short double-stranded DNA fragments with overhangs compatible with ligation into the BsmbI-digested plasmid pLKO5.sgRNA.EFS.tGFP (Addgene #57823). CRISPRi cell lines were obtained following a two-step lentiviral transduction. First, a mother cell line was created after transduction of H9 cells with the hUbC-dCas9-KRAB-T2a-Puro construct (Addgene #71236, Pontis, et al., 2019), followed by 3 days of puromycin selection at 0.2 μg/mL. A second lentiviral transduction was performed with the constructs containing the sgRNAs. The knock-down (KD) levels were assessed after a minimum of 5 days post-transduction of the guides. Multiple sgRNAs were tested, and the downstream experiments were performed with one guide, with the following targeting sequence: TACGCAATACAACTCCCGAG.

### Imaging of beating monolayers

Live imaging of the beating monolayers of cardiomyocytes was performed using Leica SP8 Inverted with the HC PL Fluotar 10x/0.3 air objective using DIC settings. The images were acquired with a Leica DFC 7000 G/T CCD camera Leica Application Suite (LAS) X software at 10 frames/sec. Time-lapse microscopy images were analyzed in python using Particle Image Velocimetry (PIV) approach described in Kumar, et al. 2019 after 5x5 binning with a window size of 32 pixels and 50% overlap, implemented *via* the OpenPIV library (Ben-Gida, et al. 2020). Velocity vectors were calculated based on the pixel displacements between consecutive frames, resulting in a set of velocity magnitudes per image acquisition. The sum of these magnitudes was used to generate a representative velocity time series for each image set. Fast Fourier transformation (FFT) was applied to these velocity time series to identify dominant beating frequencies.

### Co-immunoprecipitation

Cells were lysed in the whole cell lysis buffer (500 mM HCl pH7.4, 20 mM EDTA, 30 mM sodium pyrophosphate decahydrate, 40 mM sodium fluoride, 2 mM benzamidine, 1 mM PMSF, 0.5% IGEPAL in water with EDTA-free protease inhibitors) by incubating cells for 30 min on ice under the agitation. Lysates were sonicated for 2x5 sec and centrifugated for 10 min at 10000 rcf at 4°C. Supernatants were then quantified using BCA kit (Thermo Fisher Scientific 23227) and equal amounts of protein were incubated with anti-HA magnetic beads (Thermo Fisher Scientific, 88836), overnight on a rotating wheel at 4°C. Beads were washed 3 times in wash buffer (500 mM HCl pH7.4, 20 mM EDTA, 30 mM sodium pyrophosphate decahydrate, 40 mM sodium fluoride, 2 mM benzamidine, 0.05% IGEPAL in water with EDTA-free protease inhibitors). Immunoprecipitated proteins were collected in wash buffer supplemented with 1x protein loading dye, denatured for 10 min at 95°C and loaded onto Novex 4-20% Tris-glycine SDS-PAGE gels for protein separation. Protein transfer was done using Invitrogen iBlot3 Transfer system for 15 min at 15 V. Membranes were blocked with 5% milk in PBS/0.1% Tween for 30 min, and then incubated with anti-HA-HRP (Roche, 12013819001, 1:2000) or anti-KAP1 (Abcam, ab10483, 1:1000) in 5% milk in PBS/0.1%Tween for 1h at room temperature, washed 3 times with PBS/0.1%Tween. Anti-KAP1 membrane was further incubated with an anti-rabbit secondary antibody (Jackson ImmunoResearch 111-035-008, 1:2000) for 1h at room temperature and both membranes were incubated with the ECL substrate (Advansta K-12049-D50) for 5 min and imaged on Fusion FX Vilber imaging system (Vilber Lourmat, CH).

### RNA-extraction and RT

Total RNA-extraction was done using NucleoSpin RNA XS, Micro kit for RNA purification (Machery-Nagel) by the protocol. cDNA synthesis was done using Maxima H minus cDNA synthesis master mix (Thermo Fisher Scientific).

### RNA-sequencing

RNA-seq libraries were prepared using the Illumina Truseq Stranded mRNA kit. Libraries were sequenced in 75 or 100 bp paired-end formats on the Illumina Hiseq 4000 and NovaSeq 6000 sequencers, respectively. RNA-seq reads were mapped to the hg19 human genome assembly using hisat2 v2.2.1 (Kim, et al., 2015) with parameters hisat2 -k 5 --seed 42. Only uniquely mapped reads were used for creating gene read counts with a mapping quality over 10 (MAPQ > 10) using featureCounts v2.0.1 (Liao, et al., 2014). To avoid read assignment ambiguity between genes and TEs, a gtf file containing both was provided to featureCounts. For repetitive sequences, an in-house curated version of the Repbase database was used (fragmented LTR and internal segments belonging to a single integrant were merged). Only uniquely mapped reads were used for counting on genes and TEs. Finally, the features for which the sum of reads across all samples was lower than the number of samples were discarded from the analysis. Normalization for sequencing depth using the TMM method implemented in the limma package of Bioconductor v3.54.2 (Gentleman, et al. 2004). Counts on genes were used as the library size to correct both gene and TE expression.

Differential gene expression analysis was performed using voom (Law, et al. 2014), as implemented in the limma package of Bioconductor. A gene was considered to be differentially expressed when the fold change between groups was bigger than 2 and the p-value was smaller than 0.05. A moderated t-test (as implemented in the limma package of R) was used to test significance. P-values were corrected for multiple testing using the Benjamini-Hochberg method (Benjamini and Hochberg, 1995).

### ATAC-seq

ATAC-seq was done by the original protocol (Buenrostro, et al. 2013). Cells were detached as single cells, and each condition was processed in triplicates, as independent differentiations, with 100000 cells per replicate lysed in ATAC-seq lysis buffer (10% NP40, 10% Tween-20 and 1% digitonin in ATAC-RSD) to isolate the nuclei. After the washing, chromatin was incubated with Tn5 transposition mix (in TDE1 enzyme, 10% Tween-20, 1% digitonin in TD buffer and PBS). DNA fragments were then purified on Zymo columns (Zymo, D4014), and used as input for PCR for library amplification using custom index (Buenrostro, et al. 2013). Resulting libraries were quantified and sequenced using 75-bp paired-end sequencing on an AVITI sequencer (Element Biosciences).

Raw reads were aligned to the human genome (hg19) using Bowtie2 v2.4.5 in sensitive local mode. Bam files were filtered to remove low-quality reads (MAPQ < 10). Bigwig coverage tracks, representing the mean of the 3 replicate samples, were generated using bedtools v2.30.0. Peak calling was performed with MACS2 v2.2.7.1 using the parameters --nomodel and -- nolambda, as recommended for ATAC-seq data. Peaks were merged per condition using the bedtools merge function, and a final unified peak list was created using bedtools merge for subsequent signal quantification with bedtools genomecov. ATAC coverage signal profiles were created using the computeMatrix function and plotHeatmap from deeptools 3.5.4. ZNF436 peaks overlapping ATAC-seq peaks were used for signal quantification with UCSC tool bigWigAverageOverBed. The resulting count matrix was processed in R.

### Cleavage under targets and tagmentation (CUT&Tag)

Cleavage Under Targets and Tagmentation (CUT&Tag) was done by the original protocol (Kaya-Okur, et al. 2019). For each mark, 100k cells were used per sample with the anti-H3K9me3 primary antibody (Diagenode, C15410056), anti-H3K4me3 primary antibody (Cell Signaling Technology, C42D8, 9751S), anti-H3K4me1 primary antibody (Diagenode, C15410194), anti-H3K27ac primary antibody (Active Motif, 39034), and the anti-rabbit IgG secondary antibody (antibodies, ABIN101961). A homemade purified pA-Tn5 protein (3XFlag-pA-Tn5-Fl) was produced by the Protein Production and Purification Facility (PTPSP) at EPFL and then coupled with MEDS oligos. The purified recombinant protein was used at a final concentration of 700 ng/μL (1:250 dilution from homemade stock).

Libraries were sequenced with 75 bp paired-end reads on the NextSeq 500 (Illumina). Raw reads were mapped to the human genome (hg19) using the short-read aligner program Bowtie2 v2.4.5, with the sensitive local mode, retaining only the high-quality reads (MAPQ > 10). Bigwig coverage tracks, representing the mean of the 3 replicate samples, were generated using bedtools v2.30.0. Histone mark coverage signal profiles were created using the computeMatrix function and plotHeatmap from deeptools 3.5.4.

### Protein proximity biotinylation, pull-down and mass spectrometry (MS)

Proximity biotinylation and pull-down experiments were done by the protocol with slight modifications (Santos-Barriopedro, et al. 2021). TurboID enzyme was fused to a target protein of interest with the presence of the HA tag. The constructs were stably introduced into human ESCs using lentiviral transduction. vCMs were harvested at day 15 of differentiation. vCMs were incubated for 10 min at 37^°C^ with media supplemented with 50 uM biotin. Immediately after, cells were placed on ice, washed twice with ice-cold PBS and lysed using RIPA buffer (50 mM Tris-HCl, 150 mM NaCl, 0.1% SDS, 0.5% NaDOC, 1% NP40) with supplemented protease inhibitors (EDTA-free, Roche 11873580001). Lysate was sonicated 2x15 sec cycle for fragmenting DNA. Streptavidin beads were modified with CHD (1,2-cyclohexanedione, Sigma Aldrich, C101400-1G) and NaBH3CN (sodium cyanoborohydride, Sigma Aldrich, 8180530010) before the pull-down. For the pull-down, 300 ug of the protein was used per replicate for hESCs and 900 ug of protein for vCMs. Pull-downs were done overnight at 4° C. After three washes with RIPA, the beads with pulled-down proteome were digested at 37° C for 2h with MS-grade Lys-C (Promega, VA1170) and then for 16h with MS-grade Trypsin (Promega, V5280). After the digestion, samples were acidified using 100% formic acid and 0.1% TFA (trifluoroacetic acid). Acidified peptides were desalted in StageTips using 4 disks from an Empore SDB-RPS (3 M) filter based on the standard protocol (Rappsilber, et al., 2007). Purified peptides were dried down by vacuum centrifugation. Samples were resuspended in 2% acetonitrile (Biosolve), 0.1% FA. Nano-flow separations were performed on a Dionex Ultimate 3000 RSLC nano UPLC system (Thermo Fischer Scientific) on-line connected with an Exploris 480 Orbitrap Mass Spectrometer (Thermo Fischer Scientific). A capillary precolumn (Acclaim Pepmap C18, 3 μm-100Å, 2 cm x 75μm ID) was used for sample trapping and cleaning. A 50 cm long capillary column (75 μm ID; in-house packed using ReproSil-Pur C18-AQ 1.9 μm silica beads; Dr. Maisch) was then used for analytical separations at 250 nl/min over 150 min biphasic gradients. Acquisitions were performed through Top Speed Data-Dependent acquisition mode using a cycle time of 2 seconds. First MS scans were acquired with a resolution of 60’000 (at 200 m/z) and the most intense parent ions were selected and fragmented by High energy Collision Dissociation (HCD) with a Normalized Collision Energy (NCE) of 30% using an isolation window of 1.4 m/z. Fragmented ions were acquired with a resolution 15’000 (at 200 m/z) and selected ions were then excluded for the following 20 s.

Raw data were processed using MaxQuant 1.6.10.43 (Cox and Mann, 2008) against a database consisting of the *Homo sapiens* reference proteome database (81837 protein sequences Release 2022_05), GFP and MaxQuant contaminant protein sequences. Carbamidomethylation was set as a fixed modification, whereas oxidation (M), phosphorylation (S, T, Y), acetylation (Protein N-term) and glutamine to pyroglutamate were considered as variable modifications. A maximum of two missed cleavages were allowed and “Match between runs” option was enabled. A minimum of 2 peptides was required for protein identification and the false discovery rate (FDR) cutoff was set to 0.01 for both peptides and proteins. Label-free quantification (LFQ) was performed by MaxQuant standard pipeline and LFQ intensity counts were median normalized using the variance stabilization normalization (VSN) as implemented in the DEP R package v1.20. Missing values were imputed differently depending if the data was missing at random (MAR) or not (MNAR). For MAR, the k-nearest neighbors approach was used with option rowmax = 0.9. For MNAR values, the imputation was performed by drawing a Gaussian distribution using the minProb option with parameter q = 0.1. For the contrasts between conditions, the threshold of FDR = 0.05 and log2 fold change 1.5 were chosen.

### Immunofluorescence

Human ESCs expressing TurboID constructs were differentiated onto matrigel-coated coverslips and processed at d15 of differentiation. To initiate biotinylation, vCMs were incubated for 10 min at 37°C with media supplemented with 50 μM biotin and immediately fixed for 15 min in a 4% methanol-free formaldehyde solution. Cells were washed three times with PBS, permeabilized with 0.5 % Triton X-100 for 20 min and blocked with 1 % BSA for 30 min. Coverslips were then incubated with anti-HA-rabbit antibody (1:1000, Abcam ab9110) and streptavidin-A647 (1:1000, Invitrogen S32357) in 1 % BSA in PBS overnight at 4°C or 1h at room temperature. Samples were washed three times with 1% BSA in PBS and incubated with secondary antibody anti-rabbit-A488 (Thermo Fisher Scientific A-21206). DAPI was added in the last 10 min of secondary antibody incubation. Three final washes were performed before mounting the slides in Vectashield mounting medium (Vector Laboratories). Images were acquired on a Leica-SP8 confocal microscope using the oil objective HC PL APO 63x/1.40 and analyzed with FIJI ImageJ.

### KZFP-embedded TE stratification

KZFP ChIP-seq data was compiled from the previous studies (Schmitges, et al. 2016, Imbeault, et al. 2017, de Tribolet-Hardy, et al. 2023). Statistical enrichment of each of 301 KZFPs over TE subfamilies was performed using the pyTEnrich algorithm, which takes into account specific TE subfamily distributions and corrects for their overrepresentation (de Tribolet-Hardy, et al. 2023, https://alexdray86.github.io/pyTEnrich/build/html/index.html). If one or more KZFPs were enriched for a TE subfamily (p.adj.final < 0.05), the integrants of that subfamily were split into genomic interval files based on the KZFP binding. Otherwise, TE integrants belonging to subfamilies not enriched in the binding of any KZFP were assigned the ‘unbound’ category. KZFP-stratified TE interval files were then queried against the ENCODE data set, consisting of 6 histone marks ChIP-seqs interval files for up to 82 different tissues using GIGGLE (Layer, et al. 2018), a tool suitable for large-scale feature intersections. Thereby, each TE subfamily ended up described with a set of Giggle scores for each KZFP-stratified integrants’ overlap with tissue-specific histone marks, where a score is a composite product of mean odds ratio (MOR) and right tail of Fisher’s exact test. The matrix of GIGGLE scores for each TE subfamily was used as input for UMAP dimensionality reduction with n_neighbours = 10 (McInnes, et al. 2018).

### Computing dN/dS ratio

For the human ZNF436 gene orthologs, multiple sequence alignments were constructed using the codon_alignment.pl script (Li et al. 2017) (https://github.com/lyy005/codon_alignment/blob/master/) and used as input for the HyPhy pipeline. Specifically, we generated a phylogenetic tree of the orthologs using PhyML (Guindon et al. 2010) (v3.3.20220408). These trees were then analyzed with the HyPhy (Kosakovsky Pond, et al. 2005) (version 2.5.2) pipeline, utilizing the FEL subroutine to calculate the dN/dS ratios and to determine their significance for each nucleotide position in ZNF436 with a P value cutoff of 0.05.

### Relative tissue specificity scoring

To assess the tissue-specificity of transposable element (TE) integrants bound by the ZNF436, we analyzed H3K27ac signals across 67 human tissues from the ENCODE. First, TE integrants were treated as genomic features for which the H3K27ac signal was quantified from the read pile-ups of the ChIP-seq performed in a given tissue. Next, TE integrants were subsetted based on the overlapping ZNF436 ChIP-seq peaks, which resulted in 3711 ZNF436-bound TE integrants with detectable H3K27ac signal. Z-score normalization was applied across tissues for each TE integrant. Specificity scores were computed as the maximum absolute z-score across tissues, representing the highest relative enrichment in a single tissue. These scores were further normalized to a 0–1 scale using min-max scaling, for a relative tissue-specific score, and a threshold of the top 10% of integrants was used to define highly tissue-specific ZNF436-bound TE integrants.

### CRE activity of L2/MIR integrants

To assess the H3K27ac status of all ZNF436-bound L2/MIR integrants, we analyzed H3K27ac signals across 67 human tissues from the ENCODE. ZNF436 peaks encompassed 2884 L2/MIR integrants that were treated as genomic features for which the H3K27ac signal was quantified from the read pile-ups of the ChIP-seq performed in a given tissue, making a 2884x67 matrix used as an input for UMAP dimensionality reduction with n_neighbours = 10.

### External datasets

In this study, we utilized single-cell RNA-seq datasets from the Human Cell Atlas of fetal gene expression (Cao et al., 2020, GEO accession number: GSE156793), RNA-seq from human *in vivo* heart development (Cardoso-Moreira, et al., 2019, EMBL-EBI accession number: E-MTAB-6814), RNA-seq from human *in vitro* cardiomyocyte differentiation (Zhang, et al., 2019, GEO accession number: GSE116862), ChIP-seq for KZFP binding (Schmitges, et al., 2016, GEO accession number GSE76496; Imbeault et al., 2017, GEO accession number GSE78099; de Tribolet-Hardy et al., 2023, GEO accession number GSE20096) and ChIP-seq for histone marks in adult human tissues (The ENCODE Project Consortium, 2020; data was downloaded from the ENCODE portal https://www.encodeproject.org/ with the sample identifiers found in supplementary table 1).

### Data analysis, Statistics and Reproducibility

All data were analyzed using custom scripts executed in a bash shell on the terminal, python, and/or R. Statistical analyses were performed using R, with detailed statistical information provided in the figure legends, and associated p-values reported on the figures. Results are considered as statistically significant, with a p-value < 0.05. Significance levels were denoted as ns (not significant), *p-value < 0.05, **p-value < 0.01, ***p-value < 0.001, ****p -value < 0.0001. Data are represented as means ± standard deviations unless otherwise specified. All experiments were independently repeated at least twice with consistent results.

## Notes

### Competing Interest Statement

The authors have declared no competing interest.

## References

1. Alexander, J. M. et al. Brg1 modulates enhancer activation in mesoderm lineage commitment. Development, 142, 1418–1430 (2015). DOI: 10.1242/dev.109496

2. Andrews G. et al. Mammalian evolution of human cis-regulatory elements and transcription factor binding sites. Science, 380, 6643 (2023). DOI: 10.1126/science.abn7930

3. Appikonda S. et al. Cross-talk between chromatin acetylation and SUMOylation of tripartite motif–containing protein 24 (TRIM24) impacts cell adhesion. Journal of Biological Chemistry, 293, 19, 7476–7485 (2018). DOI: 10.1074/jbc.RA118.002233

4. Barrero, M. et al. Epigenetic Mechanisms that Regulate Cell Identity. Cell Stem Cell, 7, 5 (2010). DOI: 10.1016/j.stem.2010.10.009

5. Baxter, A. et al. The SWI/SNF chromatin remodeling complexes BAF and PBAF differentially regulate epigenetic transitions in exhausted CD8+ T cells. Immunity, 56 (2023). DOI: 10.1016/j.immuni.2023.05.008.

6. Begnis, M. et al. Clusters of lineage-specific genes are anchored by ZNF274 in repressive perinucleolar compartments. Science Advances, 10, 37 (2024). DOI: 10.1126/sciadv.ado1662

7. Ben-Gida, H. et al. OpenPIV-MATLAB — An open-source software for particle image velocimetry; test case: Birds’ aerodynamics. SoftwareX, 12 (2020). DOI: 10.1016/j.softx.2020.100585

8. Benjamini Y. and Hochberg Y. Controlling the False Discovery Rate: A Practical and Powerful Approach to Multiple Testing. Journal of the Royal Statistical Society: Series B (Methodological) (1995). DOI: 10.1111/j.2517-6161.1995.tb02031.x

9. Bourque, G. et al. Ten things you should know about transposable elements. Genome Biology, 19, 199 (2018). DOI: 10.1186/s13059-018-1577-z

10. Britten, R. J., Davidson, E. H. Repetitive and Non-Repetitive DNA Sequences and a Speculation on the Origins of Evolutionary Novelty. The Quarterly Review of Biology, 46 (1971). DOI: 10.1086/406830

11. Buenrostro, J. et al. Transposition of native chromatin for fast and sensitive epigenomic profiling of open chromatin, DNA-binding proteins and nucleosome position. Nature Methods, 10, 1213–1218 (2013). DOI: 10.1038/nmeth.2688

12. Cardoso-Moreira, M. et al. Gene expression across mammalian organ development. Nature, 571, 505–509 (2019). DOI: 10.1038/s41586-019-1338-5

13. Cao, J. et al. A human cell atlas of fetal gene expression. Science, 370, 6518 (2020). DOI: 10.1126/science.aba7721

14. Cao, Y. et al. Widespread roles of enhancer-like transposable elements in cell identity and long-range genomic interactions. Genome Research, 29, 49–52 (2019). DOI: 10.1101/gr.235747.118

15. Carroll, S. B. Evo-Devo and an Expanding Evolutionary Synthesis: A Genetic Theory of Morphological Evolution. Cell, 134 (2008). DOI: 10.1016/j.cell.2008.06.030

16. Chuong, E. et al. Regulatory activities of transposable elements: from conflicts to benefits. Nature Reviews Genetics 18, 86 (2017). DOI: 10.1038/nrg.2016.139

17. Cox, J. and Mann, M. MaxQuant enables high peptide identification rates, individualized p.p.b.-range mass accuracies and proteome-wide protein quantification. Nature Biotechnology 26, 1367–1372 (2008). DOI: 10.1038/nbt.1511

18. De Franco, E. et al. Primate-specific ZNF808 is essential for pancreatic development in humans. Nature Genetics 55, (2023). DOI: 10.1038/s41588-023-01565-x

19. de Tribolet-Hardy, J. et al. Genetic features and genomic targets of human KRAB-zinc finger proteins. Genome Research, 33, 1409–1423 (2023). DOI: 10.1101/gr.277722.123

20. Deininger, P. Alu elements: know the SINEs. Genome Biology 12, 236 (2011). DOI: 10.1186/gb-2011-12-12-236

21. Domcke, S. et al. A human cell atlas of fetal chromatin accessibility. Science, 370, 6518 (2020). DOI: 10.1126/science.aba7612

22. Du, A.Y. et al. Regulatory transposable elements in the encyclopedia of DNA elements. Nature Communications, 15, 7594 (2024). DOI: 10.1038/s41467-024-51921-6

23. Emerson, R. O. and Thomas, J. H. Gypsy and the Birth of the SCAN Domain. Journal of Virology, (2011). DOI: 10.1128/jvi.00867

24. Fillot, T. and Mazza, D. Rethinking chromatin accessibility: from compaction to dynamic interactions. Current Opinion in Genetics & Development, 90 (2025). DOI: 10.1016/j.gde.2024.102299

25. Forey, et al. Cancer cells subvert the primate-specific KRAB zinc finger protein ZNF93 to control APOBEC3B. bioRxiv (2025). DOI: 10.1101/2025.03.05.641617

26. Gassler, J. et al. Zygotic genome activation by the totipotency pioneer factor Nr5a2. Science, 378, 6626, 1305–1315 (2022). DOI: 10.1126/science.abn7478

27. Gentleman, R. C. et al. Bioconductor: open software development for computational biology and bioinformatics. Genome Biology, 5 (2004). DOI: 10.1186/gb-2004-5-10-r80

28. Grossi, E. et al. The SWI/SNF PBAF complex facilitates REST occupancy at repressive chromatin. Molecular Cell, 85 (2025). DOI: 10.1016/j.molcel.2025.03.026

29. Guindon, S. et al. New Algorithms and Methods to Estimate Maximum-Likelihood Phylogenies: Assessing the Performance of PhyML 3.0. Systematic Biology, 59, 3, (2010). DOI: 10.1093/sysbio/syq010

30. Helleboid, P-Y. et al. The interactome of KRAB zinc finger proteins reveals the evolutionary history of their functional diversification. The EMBO Journal, 38 (2019). DOI: 10.15252/embj.2018101220

31. Hota, S. K. et al. Dynamic BAF chromatin remodeling complex subunit inclusion promotes temporally distinct gene expression programs in cardiogenesis. Development, 146, 19 (2019). DOI: 10.1242/dev.174086

32. Hota, S. K. et al. Brahma safeguards canalization of cardiac mesoderm differentiation. Nature 602, 129–134 (2022). DOI: 10.1038/s41586-021-04336-y

33. Hyacinthe, J. and Bourque, G. Transposable elements impact the regulatory landscape through cell type specific epigenomic associations. Biorxiv (2024). DOI: 10.1101/2024.08.07.606967

34. Imbeault, M. et al. KRAB zinc-finger proteins contribute to the evolution of gene regulatory networks. Nature, 543, 550–554 (2017). DOI: 10.1038/nature21683

35. Jjingo, D. et al. Mammalian-wide interspersed repeat (MIR)-derived enhancers and the regulation of human gene expression. Mobile DNA 5, 14 (2014). DOI: 10.1186/1759-8753-5-14

36. Isbel, L. et al. Readout of histone methylation by Trim24 locally restricts chromatin opening by p53. Nature Structural & Molecular Biology, 30, 948–957 (2023). DOI: 10.1038/s41594-023-01021-8

37. Iouranova, A. et al. KRAB zinc finger protein ZNF676 controls the transcriptional influence of LTR12-related endogenous retrovirus sequences. Mobile DNA 13, 4 (2022). DOI : 10.1186/s13100-021-00260-0

38. Kaya-Okur, H. S., et al. CUT&Tag for efficient epigenomic profiling of small samples and single cells. Nature Communications, 10, 1930 (2019). DOI: 10.1038/s41467-019-09982-5

39. Kim, D. et al. HISAT: a fast spliced aligner with low memory requirements. Nature Methods, 12, 4, 357–360 (2015). DOI: 10.1038/nmeth.3317

40. Klemm, S.L. et al. Chromatin accessibility and the regulatory epigenome. Nature Reviews Genetics, 20, 207–220 (2019). DOI: 10.1038/s41576-018-0089-8

41. Kramerov, D. and Vassetzky, N. Origin and evolution of SINEs in eukaryotic genomes. Heredity 107, 487–495 (2011). DOI: 10.1038/hdy.2011.43

42. Kosakovsky Pond S. L. et al. HyPhy: hypothesis testing using phylogenies, Bioinformatics, 21, 5 (2005). DOI: 10.1093/bioinformatics/bti079

43. Kumar, N. et al. Assessment of temporal functional changes and miRNA profiling of human iPSC-derived cardiomyocytes. Scientific Reports, 9, 13188 (2019). DOI: 10.1038/s41598-019-49653-5

44. Lambard, S. et al. Expression of rod-derived cone viability factor: dual role of CRX in regulating promoter activity and cell-type specificity. PLoS One, 5, 10 (2010). DOI: 10.1371/journal.pone.0013075.

45. Lander, E. S. et al. Initial sequencing and analysis of the human genome. Nature 409, 860–921 (2001). DOI: 10.1038/35057062

46. Layer, R. M. et al. GIGGLE: a search engine for large-scale integrated genome analysis. Nature Methods, 15, 123–126 (2018). DOI: 10.1038/nmeth.4556

47. Law, C.W. et al. voom: precision weights unlock linear model analysis tools for RNA-seq read counts. Genome Biology, 15, (2014). DOI: 10.1186/gb-2014-15-2-r29

48. Li, Y. et al. The molecular evolutionary dynamics of oxidative phosphorylation (OXPHOS) genes in Hymenoptera. BMC Evolutionory Biology, 17, 269 (2017). DOI: 10.1186/s12862-017-1111-z

49. Liang, Q. et al. Linking a cell-division gene and a suicide gene to define and improve cell therapy safety. Nature 563, 701–704 (2018). DOI: 10.1038/s41586-018-0733-7

50. Liao, Y. et al. featureCounts: an efficient general purpose program for assigning sequence reads to genomic features. Bioinformatics, 30, 7, 923–930, (2014). DOI: 10.1093/bioinformatics/btt656

51. Matsushima, W. et al. Ancestral genome reconstruction enhances transposable element annotation by identifying degenerate integrants. Cell Genomics 4, 2 (2024). DOI: 10.1016/j.xgen.2024.100497

52. Matsushima, et al. Zinc-finger proteins with a co-opted capsid domain anchor nucleosomes over transposon sequences. BioRxiv (2025). DOI: 10.1101/2025.03.03.638093

53. McInnes, L. et al. UMAP: Uniform Manifold Approximation and Projection. The Journal of Open Source Software, 3, 29, 861 (2018). DOI: 10.21105/joss.00861

54. Mills, R. et al. Which transposable elements are active in the human genome? Trends in Genetics, Trends in Genetics, 23, 4 (2007). DOI:

55. Moore, D., et al. The KRAB-Zinc Finger protein ZKSCAN3 represses enhancers via embedded retrotransposons. bioRxiv (2025). DOI: 10.1101/2025.01.30.635440

56. Ohno, S. Evolution by Gene Duplication (Springer, New York, 1970). DOI: 10.1007/978-3-642-86659-3

57. Orgel, L. E., Crick F. H. C. Selfish DNA: the ultimate parasite. Nature 284, 604–607 (1980). DOI: 10.1038/284604a0

58. Osmanski, A. et al. Insights into mammalian TE diversity through the curation of 248 genome assemblies. Science, 380, 6643 (2023). DOI: 10.1126/science.abn1430

59. Plaisance, I. et al. A transposable element into the human long noncoding RNA CARMEN is a switch for cardiac precursor cell specification. Cardiovascular Research, 119, 6 (2023). DOI: 10.1093/cvr/cvac191

60. Playfoot C. J. et al. Transposable elements and their KZFP controllers are drivers of transcriptional innovation in the developing human brain. Genome Research, 31: 1531–1545 (2021). DOI: 10.1101/gr.275133.120

61. Pontis, J. et al. Hominoid-Specific Transposable Elements and KZFPs Facilitate Human Embryonic Genome Activation and Control Transcription in Naive Human ESCs. Cell Stem Cell, 24, 5, 724–735 (2019). DOI: 10.1016/j.stem.2019.03.012

62. Pontis, J., Pulver, C., Playfoot, C.J. et al. Primate-specific transposable elements shape transcriptional networks during human development. Nature Communications 13, 7178 (2022). DOI: 10.1038/s41467-022-34800-w

63. Pulver, C. et al. Statistical learning quantifies transposable element-mediated cis-regulation. Genome Biology 24, 258 (2023). DOI: 10.1186/s13059-023-03085-7

64. Rappsilber, J., Mann, M. & Ishihama, Y. Protocol for micro-purification, enrichment, pre-fractionation and storage of peptides for proteomics using StageTips. Nature Protocols 2, (2007). DOI: 10.1038/nprot.2007.261

65. Roller, M. et al. LINE retrotransposons characterize mammalian tissue-specific and evolutionarily dynamic regulatory regions. Genome Biology, 22, 62 (2021). DOI: 0.1186/s13059-021-02260-y

66. Rosspopoff, O. and Trono, D. Take a walk on the KRAB side. Trends in Genetics, 39, 11, 844–857 (2023). DOI: 10.1016/j.tig.2023.08.003

67. Rosspopoff, O., et al. Transposable element co-option drives transcription factor neofunctionalization. BiorXiv (2025). DOI: 10.1101/2025.03.01.640934

68. Rowe, H. et al. TRIM28 repression of retrotransposon-based enhancers is necessary to preserve transcriptional dynamics in embryonic stem cells. Genome Research, 23 (2012). DOI: 10.1101/gr.147678.112

69. Santos-Barriopedro, I. et al. Off-the-shelf proximity biotinylation for interaction proteomics. Nature Communications, 12, 5015 (2021). DOI: 10.1038/s41467-021-25338-4

70. Schmitges, F. W. et al. Multiparameter functional diversity of human C2H2 zinc finger proteins. Genome Research 26: 742–1752 (2016). DOI: 10.1101/gr.209643.116

71. Sekirnik A. R. et al. Identification of Histone Peptide Binding Specificity and Small-Molecule Ligands for the TRIM33α and TRIM33β Bromodomains. ACS Chemical Biology, 17, 10 (2022). DOI: 10.1021/acschembio.2c00266

72. Soleimani, V. et al. REST/NRSF preserves muscle stem cell identity and survival by repressing alternate cell fates. Research Square (2024) DOI: 10.21203/rs.3.rs-4396883/v1

73. Stoll, G. A. et al. Structure of KAP1 tripartite motif identifies molecular interfaces required for retroelement silencing. Proceedings of the National Academy of Sciences, 116, 15042–15051 (2019). DOI: 10.1073/pnas.1901318116

74. The ENCODE Project Consortium, et al. Expanded encyclopaedias of DNA elements in the human and mouse genomes. Nature, 583, 699–710 (2020). DOI: 10.1038/s41586-020-2493-4

75. Tycko, J. et al. High-Throughput Discovery and Characterization of Human Transcriptional Effectors. Cell, 183, 7 (2020). DOI: 10.1016/j.cell.2020.11.024

76. Visel A. et al. VISTA Enhancer Browser-- a database of tissue-specific human enhancers. Nucleic Acid Research, 3, D88– D92 (2007). DOI: 10.1093/nar/gkl822

77. Wolf, G. et al. KRAB-zinc finger protein gene expansion in response to active retrotransposons in the murine lineage. eLife, 9 (2020). DOI: 10.7554/eLife.56337

78. Yang, P. et al. The Role of KRAB-ZFPs in Transposable Element Repression and Mammalian Evolution. Trends in Genetics, 33, 11 (2017). DOI: 10.1016/j.tig.2017.08.006

79. Zhang, Y. et al. Transcriptionally active HERV-H retrotransposons demarcate topologically associating domains in human pluripotent stem cells. Nature Genetics 51, 1380–1388 (2019). DOI: 10.1038/s41588-019-0479-7

80. Zuckerkandl, E., and Pauling, L. (1965). Evolutionary divergence and convergence in proteins. Evolving Genes and Proteins, 97-166 (1965). DOI: 10.1016/B978-1-4832-2734-4.50017-6

81. https://alexdray86.github.io/pyTEnrich/build/html/index.html

82. https://www.encodeproject.org/

83. https://portals.broadinstitute.org/gppx/crispick/public

84. https://portals.broadinstitute.org/gpp/public/resources/protocols

